# Pancreatic Brsk2 amplifies parasympathetic signals to promote type 2 diabetes

**DOI:** 10.1101/2021.08.05.455219

**Authors:** Rufeng Xu, Kaiyuan Wang, Zhengjian Yao, Li Jin, Jing Pang, Yuncai Zhou, Kai Wang, Dechen Liu, Yaqin Zhang, Peng Sun, Fuqiang Wang, Xiaoai Chang, Yating Li, Shusen Wang, Yalin Zhang, Shuyong Lin, Cheng Hu, Yunxia Zhu, Xiao Han

## Abstract

The parasympathetic nervous system (PNS) modulates postprandial glucose metabolism via innervating pancreas; however, its significance in the pathogenesis of type 2 diabetes (T2DM) remains unclear. Here we show that brain-specific serine/threonine-protein kinase 2 (Brsk2), accumulated in obese mouse islets, responds to PNS activation and initiates pre-absorptive insulin release. In inducible mouse models, excessive Brsk2 amplifies parasympathetic signaling to β cells and increases their secretion, ensuing insulin resistance and T2DM. Conversely, Brsk2 inhibition prevents and treats HFD-induced metabolic abnormities via avoiding β-cell oversecretion. Mechanistically, parasympathetic acetylcholine activates cholinergic receptor M3 (Chrm3), then Chrm3 recruits and stabilizes Brsk2, which in turn phosphorylates phospholipase A2 activating protein (Plaa). A Chrm3-Brsk2-Plaa axis stimulates β-cell hypersecretion during both pre-absorptive and absorptive stages in HFD-feeding mice, thus imposing insulin resistance and β-cell dysfunction. Blocking parasympathetic signaling to β cells by Brsk2 protein restoration, autonomic mediation drugs, or vagotomy restricted diabetes development. Moreover, three human BRSK2 variants are associated with hyperinsulinemia, insulin resistance, and T2DM in the Chinese population. These findings reveal that Brsk2 links parasympathetic nervous system to nutrition-overload induced T2DM.

## INTRODUCTION

The epidemic of obesity contributes to the burden of type 2 diabetes mellitus (T2DM) worldwide (Franks & McCarthy, 2016). Key features of obesity-associated diabetes are hyperinsulinemia and hyperglycemia due to insulin resistance and pancreatic β-cell dysfunction (Kahn, Hull et al., 2006). In general, glycemic homeostasis is achieved by a balance between glucose production in the liver and peripheral glucose metabolism. This balance depends on a series of processes regulated by insulin and glucagon secretion from islet β and α cells, coupled with the feeding and fast status (Gromada, Chabosseau et al., 2018). Hormonal inabilities to suppress hepatic glucose production (HGP) or to store energy postprandially as fat in adipose tissues lead to hyperglycemia (Roden & Shulman, 2019). During T2DM disease progression, obesity and insulin resistance increase insulin secretion to trigger a vicious cycle of hyperinsulinemia and insulin resistance that ultimately results in β-cell failure (Remedi & Emfinger, 2016). The cause-and-effect relationship between hyperinsulinemia and insulin resistance remains unresolved and hotly debated, while preclinical evidence speculates on mild suppression of hyperinsulinemia to treat obesity and insulin resistance (Page & Johnson, 2018). Identification of the originating factors that trigger the development of hyperinsulinemia independent of insulin resistance may help to unmask the dilemma and further shed light on mining approaches for clinical translation.

Many insulinotropic factors, including free fatty acids (FFAs) and GLP-1 promote insulin secretion by activating different G-protein-coupled receptors (GPCRs) through Gq and Gs mediated mechanisms that are either dependent or independent of the blood glucose level (Winzell & Ahren, 2007). Parasympathetic nervous system (PNS), engaging both Gq and Gs pathways plays an important role on glucose metabolism. Physiologically, parasympathetic acetylcholine triggered by food intakes initiates preabsorptive or cephalic phase insulin secretion to control postprandial glucose homeostasis (Teff, 2011), including synchronization of the islets to allow oscillations of islet-hormone secretion, and optimization of islet-hormone secretion under hypoglycemia condition (Ahren, 2000, Cryer, 2004, Cryer, 2006). However, denervation of the human pancreas by vagotomy or medication barely affects glucose homeostasis (Boyle, Liggett et al., 1988, Corrall & Frier, 1981), suggesting that this islet innervation is likely an adaptive measure that may be associated with islet adaptation to insulin resistance and T2DM. Indeed, human pancreases infiltrated with fat display a remodeling of islet autonomic innervation (Tang, Baeyens et al., 2018), and alterations in cellular neurotransmitter receptors result in β-cell dysfunction and human diabetes (Gautam, Han et al., 2006, Rosengren, Jokubka et al., 2010). Interestingly, rodents fed an HFD for 8 weeks increase carbachol, an acetylcholine analog-stimulated insulin secretion (Ahren, Simonsson et al., 1997), strengthening the notion that PNS activation imposes obesity-induced hyperinsulinemia (Magnan, Collins et al., 1999). However, the molecules that coordinate the PNS signals in islet hormonal cells remain unclear.

One possible candidate is brain-specific serine/threonine-protein kinase 2 (*Brsk2*), a member of the AMP-activated protein kinase (AMPK)-related kinase family. Brsk2, selectively expressed in the pancreas and brain, defines neuronal polarization and forms central axon arbors for sensory neurons (Nie, Lilley et al., 2013b). Mice with global or pancreas-specific Brsk2 deletion remain healthy and fertile, but show decreased serum insulin levels (Nie, Liu et al., 2013c, Nie, Sun et al., 2012), suggesting that Brsk2 is important for β-cell biology. Previously, we have showed that Brsk2 as a down-stream molecule of mTOR/S6K could trigger massive insulin secretion and enlarge β-cell size in in vitro model (Nie et al., 2013b, Nie et al., 2012). However, the in vivo role of Brsk2 during the pathogenesis of T2DM is still poorly understood. The aim of the present study was to assess whether Brsk2 integrates autonomic, nutritional, and gut hormonal signals to regulate β-cell function and the development of T2DM by using inducible β-cell specific Brsk2 gain-of-function and loss-of-function mouse models. Our findings indicate that β-cell Brsk2 links PNS to T2DM in the context of metabolic stress. Notably, human BRSK2 variants are associated with glucose metabolism. Pharmacological inhibition of Brsk2 improves β-cell function and glucose metabolism in HFD-fed mice, therefore BRSK2 may be a promising therapeutic target for restricting obesity-induced hyperinsulinemia, thus preventing and treating T2DM.

## RESULTS

### Obesity-induced Brsk2 elevation connects to hyperinsulinemia via Gq-but not Gs-coupled pathways

Hyperinsulinemia is the primary cause of obesity, which ensues insulin resistance and T2DM. Our previous studies have shown that Brsk2 positively regulates insulin secretion (Nie et al., 2013b, Nie et al., 2013c). Here we investigated the relationship between obesity and β-cell Brsk2 level by using HFD mouse model. Compared with mice fed a normal chow-diet (NCD), HFD-fed mice showed significant increases in body weights from 4 to 16 weeks. Most of the mice manifested T2DM after 16 weeks of HFD feeding (Supporting Information Fig S1 A to E), and their Brsk2 protein expression was increased after 4-weeks HFD feeding (Fig 1A). However, the Brsk2 mRNA level was not altered (Supporting Information Fig S1F).

**Figure 1.**
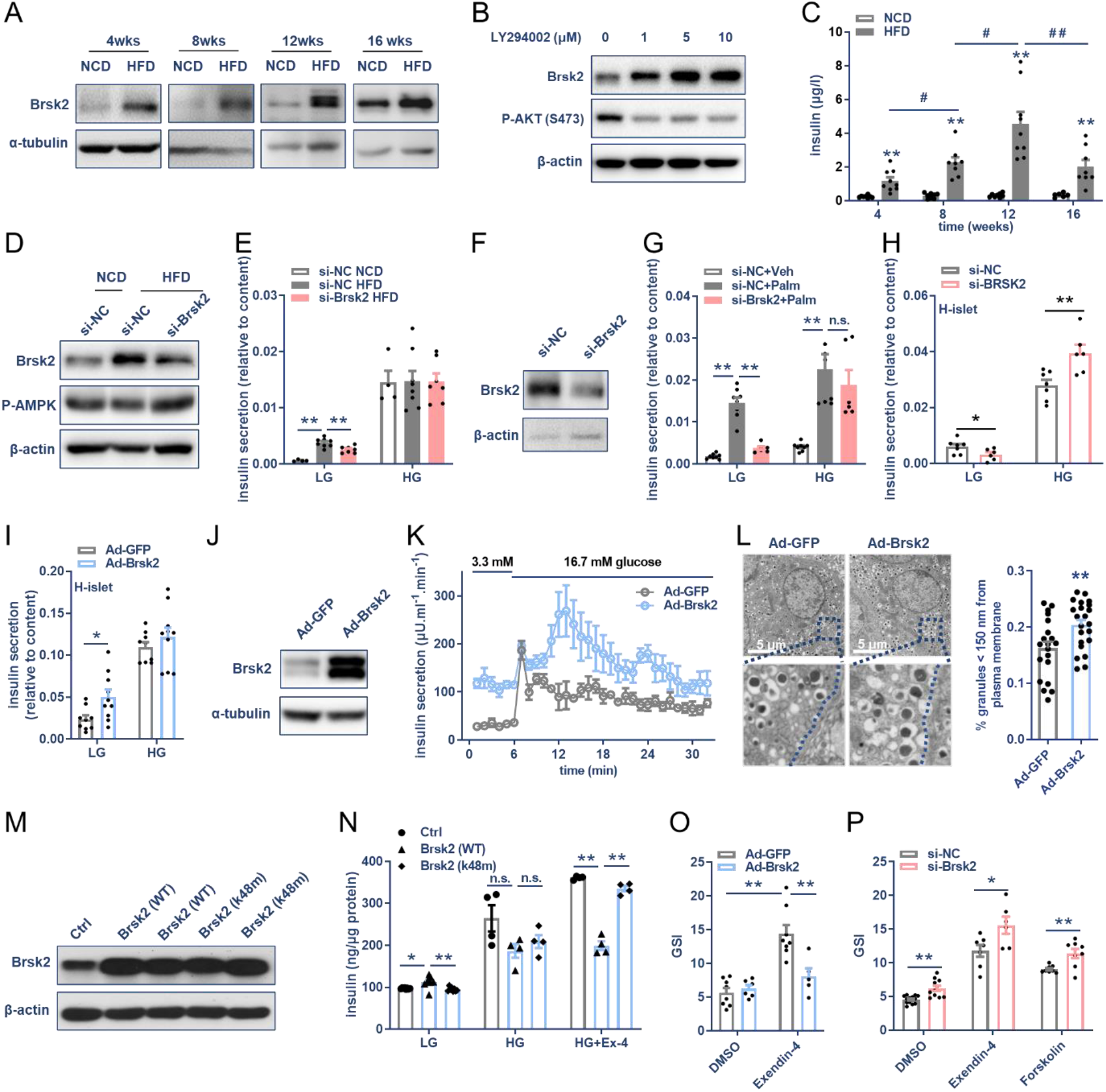
Obesity-induced Brsk2 elevation connects to hyperinsulinemia via Gq-but not Gs-coupled pathways. (A) Immunoblots showing the protein levels of Brsk2 in primary islets from mice fed on NCD or HFD for indicated weeks. (B) MIN6 cells were treated with LY294002 at indicated dose for 24 h, and protein levels were detected. (C) Serum insulin levels of mice with NCD or HFD feeding for indicated weeks. (D,E) Eight-week HFD islets were transfected with si-Brsk2 or si-NC for 48 h, and Brsk2 interference efficiency was detected (D). Then GSIS assays were performed (E). LG = 3.3 mmol/l; HG = 16.7 mmol/l glucose. (F,G) Mouse islets transfected with si-Brsk2 for 48 h for protein detection, and GSIS assays with palmitate (Palm, 0.4 mM) or vehicle (Veh) for 2 h. (H,I) Human islets were transfected with si-Brsk2 for 48 h (H) or infected with Ad-Brsk2 for 24 h (I), and GSIS assays were performed. (J,K) Mouse islets were infected with Ad-Brsk2 for 24 h for protein examination and islet perfusion. (L) Ad-Brsk2 infected mouse islets were analyzed by transmission electron microscopy. Granules nearby the plasma membrane less than 150 nm were counted to show membrane fused insulin granules. Ad-GFP: n = 20; Ad-*Brsk2*: n = 22. (M,N) Stable MIN6 cell lines overexpressed with Brsk2 wide-type (WT) or kinase-dead mutant (k48m) were confirmed by immunoblots and subjected to GSIS and glucose+Exendin-4 (HG+Ex-4)-stimulated insulin secretion assays. (O,P) Mouse islets were infected with Ad-*Brsk2* for 24 h or transfected with si-*Brsk2* for 48 h and GSIS assays were performed. Data presented as Mean ± SEM. n = 3-10 per group. \**P <* 0.05, \*\**P <* 0.01.

C-terminal of Brsk2 contains multiple autoinhibitory phosphorylation sites (Lilley, Pan et al., 2013). Treatment with dephosphorylation enzyme CIAP dose-dependently decreased Brsk2 protein level (Supporting Information Fig S1G), while deleting Brsk2 C-terminal significantly shortened its protein half-life to 3.5 h (Supporting Information Fig S1 H and I). According to NetPhos software, GSK3, cdk5, PKC and p38 are top four kinases potentially phosphorylating Brsk2 (Supporting Information Fig S1J). Inhibitors of GSK3, cdk5 and PKC decreased Brsk2 protein levels (Supporting Information Fig S1K), while p38 inhibitor had negligible effect (Supporting Information Fig S1L). Conversely, inhibition of AKT significantly increased Brsk2 amount (Fig 1B). Above data implicate that phosphorylation of autoinhibitory sites stabilize Brsk2 protein, and insulin resistance may engage this to increase Brsk2 level.

Hyperinsulinemia and hyperproinsulinemia developed concurrently with increased Brsk2 (Fig 1C and Supporting Information Fig S1M). HFD islets with Brsk2 knockdown significantly reduced basal insulin secretion, thereby preserving their glucose-stimulated index (GSI) (Fig 1, D and E and Supporting Information Fig S1N). Brsk2 knockdown also inhibited palmitate-potentiated basal insulin secretion and rescued GSIS function in mouse islets (Fig 1 F and G and Supporting Information Fig S1O). Inhibition of Brsk2 also preserved GSIS function in human islets (Fig 1H and Supporting Information Fig S1P).

On the contrary, overexpression of Brsk2 in human islets increased basal-but not glucose-stimulated insulin secretion, thus exhibiting an impaired GSIS function (Fig 1I and Supporting Information Fig S1 Q to S). Mouse islet perfusion assays showed that Brsk2 overexpression increased both basal-and glucose-stimulated insulin secretion (Fig 1 J and K), however, a decreased GSI value (Supporting Information Fig S1T). Increased membrane docking insulin granules supported an augmentation of basal insulin secretion by Brsk2 (Fig 1L). Therefore, both human and mouse islets overexpressed with Brsk2 essentially phenocopied the GSIS defect of HFD mice.

Transient overexpression of Brsk2 either potentiated or suppressed Glp-1R agonist Exendin-4-stimulated insulin secretion (Guo, Tang et al., 2006, Nie et al., 2013b), which might cause undesirable results. Therefore, we constructed stably overexpressing wild type (WT) or kinase-dead mutant (k48m) Brsk2 MIN6 cell lines to overcome those limitations (Fig 1M). WT-Brsk2 cells increased basal insulin secretion compared with both control and k48m-Brsk2 cells. However, Exendin-4-potentiated insulin secretion was missing in WT-Brsk2 cells (Fig 1N), as well as in mouse islets infected with Ad-Brsk2 (Fig 1O). Conversely, interfering Brsk2 significantly enhanced Exendin-4-and Forskolin-potentiated insulin secretion (Fig 1P). These findings indicate that obesity-induced Brsk2 elevation mainly increases basal insulin secretion, which may contribute to hyperinsulinemia in obesity and prediabetic stage via Gq-coupled independent of Gs-coupled pathways.

### Hyperinsulinemia is directly modulated by Brsk2 elevation in β cells

We examined the in vivo role of Brsk2 by generating Brsk2 knockout (βKO) mice with tamoxifen-induction system (Fig 2A). The Brsk2 expression at both the mRNA and protein levels was greatly decreased in βKO islets without affecting other tissues (Fig 2 B and C, and Supporting Information Supporting Information Fig S2A). Moreover, HFD feeding no longer enhanced Brsk2 amount in βKO islets (Fig 2D).

**Figure 2.**
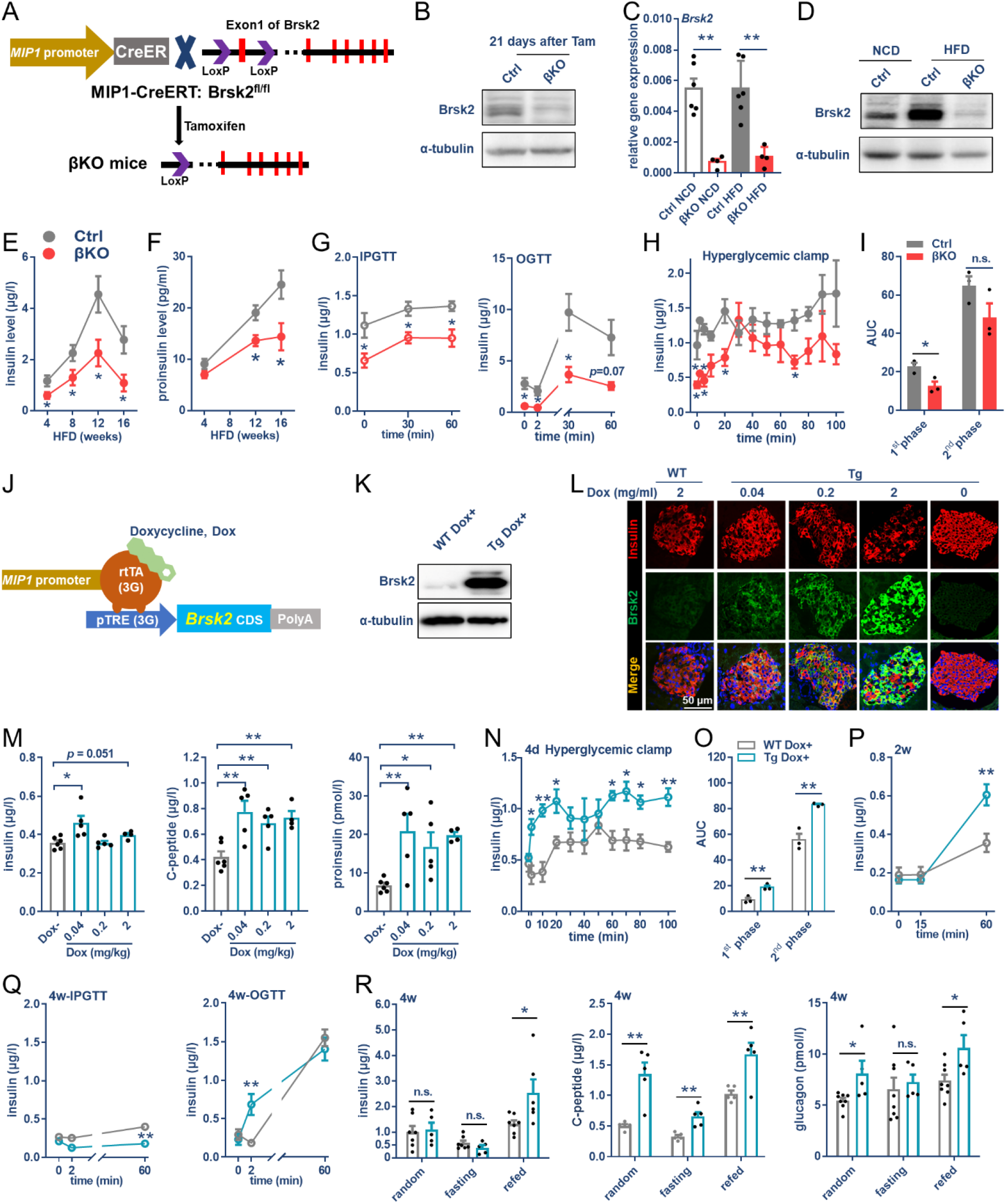
Hyperinsulinemia is directly modulated by Brsk2 elevation in β cells. (A) Schematic of β-cell specific *Brsk2* knockout (βKO) mice. (B) Protein levels of Brsk2 in βKO and Ctrl mouse islets post tamoxifen injection for 21 days. (C,D) mRNA and protein levels of Brsk2 in islets from βKO and Ctrl mice fed on HFD for 12 weeks. (E,F) Serum levels of insulin and proinsulin in βKO and Ctrl mice. (G to I) Serum insulin levels via IPGTT and OGTT (G) or during hyperglycemia-clamps (H) of βKO and Ctrl mice fed with HFD for 12 weeks. Area under the curve (AUC) of insulin secretion at first phase (1^st^ phase) and second phase (2^nd^ phase) were examined (I). (J) Schematic of β-cell specific *Brsk2* transgenic mice (MIP1-rtTA; *Brsk2*), using Tet-On system. (K) Brsk2 protein levels in islets from WT and Tg mice drinking with 2 mg/ml Dox-water for 7 days. (L) Immunofluorescence staining of Brsk2 (green), insulin (red) and nuclei (blue) in pancreas slices from WT and Tg mice drinking with indicated concentrations of Dox-water for 4 days. (M) Serum levels of insulin, C-peptide, and proinsulin were detected in Tg and WT mice with Dox induction for 4 days. (N,O) Serum insulin levels during hyperglycemia-clamps (N). AUCs of insulin secretion at 1^st^ and 2^nd^ phase (O). (P,Q) Serum insulin levels were detected via IPGTT assays after 2w-Dox induction (P), or via IPGTT and OGTT assays after 4w-Dox induction (Q). (R) Serum levels of insulin, C-peptide and glucagon at random, fasting and refed were examined after 4w-Dox induction. Data were presented as Mean ± SEM. n = 4-15 per group. \**P <* 0.05, \*\**P <* 0.01.

βKO mice fed the NCD showed normal serum insulin level, β-cell number, and islet mass (data not shown), inconsistent with global-and pancreas-Brsk2 knockout mice (Nie et al., 2013c). Notably, the βKO mice were resistant to HFD-induced hyperinsulinemia and hyperproinsulinemia (Fig 2 E and F). βKO mice limited insulin secretion upon both intraperitoneal (IPGTT) and oral (OGTTs) glucose tolerant tests (Fig 2G). Clamping the blood glucose level at ∼200 mg/dL, the glucose infusion rates (GIRs) were comparable between controls and βKO mice (Supporting Information Supporting Information Fig S2 B and C). When control mice barely responded to persistent hyperglycemia, βKO mice still secreted insulin in a biphasic manner (Fig 2 H and I).

The possibility that overexpression of Brsk2 would recall HFD-induced hyperinsulinemia was examined by generating doxycycline (Dox)-inducible, MIP1-controlled β-cell-specific Brsk2 transgenic (Tg) mice using Tet-On system (MIP1-rtTA: Brsk2; Fig 2J), thereby avoiding developmental deficiency. Mice harboring three different copy numbers of MIP1-rtTA: Brsk2 were shown as follows: Line20 (13 copies), Line42 (35 copies), and Line29 (42 copies) (Supporting Information Supporting Information Fig S2D). Administration of Dox increased islet Brsk2 levels in a copy number–dependent manner, but not in other tissues (Supporting Information Supporting Information Fig S2E). Immunostaining confirmed that Brsk2 overexpression was restricted to β cells and lowered insulin staining (Supporting Information Supporting Information Fig S2F). All three lines showed increased serum levels of C-peptide and proinsulin while slightly decreased insulin level in Line42 and Line29 mice, probably due to their reduced islet mass (Supporting Information Supporting Information Fig S2 G and H), suggesting that Brsk2 overexpression might cause persistent β-cell exocytosis. No alterations of islet mass were observed in Line20 mice, therefore Line20 mice with Dox administration (Tg Dox+) and WT Dox+ controls were used in most assays.

Again, Brsk2 levels of Tg Dox+ mice were increased in a Dox-dose dependently manner (Fig 2 K and L). Serum levels of insulin, C-peptide and proinsulin were significantly increased at 0.04 mg/ml Dox (Fig 2M). Glucose infusion provoked more insulin secretion in the Tg Dox+ mice than in the WT Dox+ mice (Fig 2N), while the GIR was comparable between them (Supporting Information Supporting Information Fig S2, I and J). The calculated first-and second-phase of insulin amounts were significantly higher in the Tg Dox+ mice (Fig 2O), akin to that of HFD mice via hyperglycemia-clamps (Supporting Information Supporting Information Fig S2 K to N). IPGTT still provoked more insulin in Tg Dox+ mice with 2-week Dox-induction (Fig 2P). However, IPGTT could not stimulate insulin in the Tg Dox+ mice after 4 weeks, by which OGTT still stimulated more insulin before glucose absorption (Fig 2Q), akin to the phenomena observed in 12-week HFD-fed mice (Supporting Information Supporting Information Fig S2 O to R). Accordingly, feeding triggered higher levels of serum insulin, C-peptide and glucagon in the Tg Dox+ mice (Fig 2R), suggesting the presence of insulinotropic stimuli apart from blood glucose level.

Taken together, by using β-cell specific gain-of-function and loss-of-function mouse models, we demonstrate that HFD-induced hyperinsulinemia relies on Brsk2 elevation at both nutrient preabsorbed and absorbed stages.

### Insulin resistance and diabetes are secondary to hyperinsulinemia caused by Brsk2 elevation

Hyperinsulinemia may result in obesity and insulin resistance. Therefore, body weights, insulin tolerance tests (ITTs) and blood glucose levels were monitored in Tg Dox+ and WT Dox+ mice after hyperinsulinemia. Tg Dox+ mice showed mild and severe insulin resistance at 2-week and 4-week Dox-inductions respectively (Fig 3A). Correspondingly, Tg Dox+ mice only showed refed hyperglycemia initially, and displayed fasting and refed hyperglycemia by 4 weeks post Dox-induction (Fig 3B). After that, the Tg Dox+ mice showed appreciable weight loss by day 37 (Supporting Information Supporting Information Fig S3A); therefore, a metabolic cage was introduced. Daily food and drinking water intakes were significantly increased in the Tg Dox+ mice; however, their feeding behaviors were not affected (Supporting Information Supporting Information Fig S3B). The Tg Dox+ mice favored lipids over glucose as an energy source, judging by the decreased respiratory exchange rate (RER) (Supporting Information Supporting Information Fig S3B). These data suggest that mice with prolonged β-cell Brsk2 overexpression manifests advanced diabetes symptoms.

**Figure 3.**
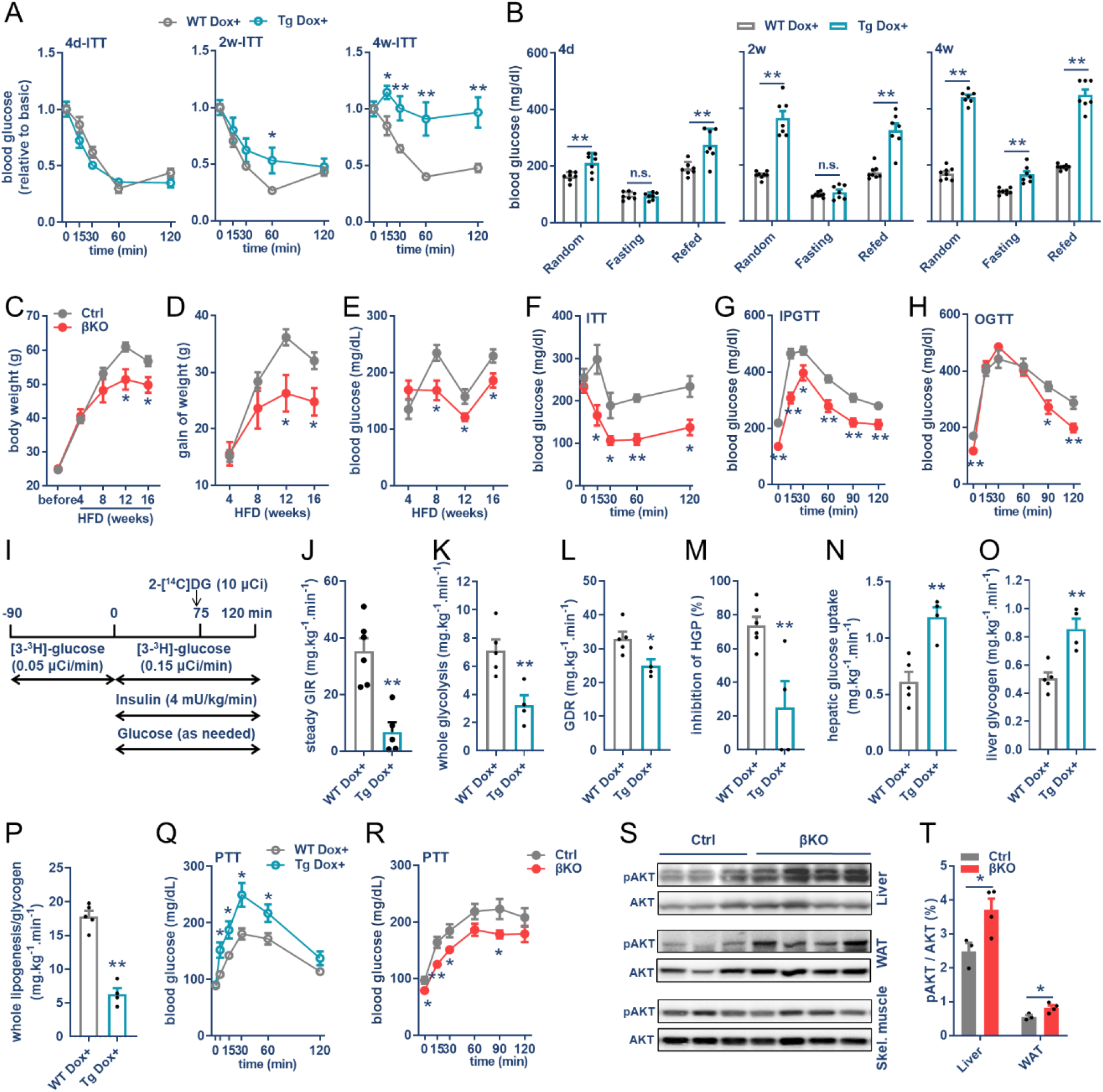
Insulin resistance and diabetes are secondary to hyperinsulinemia caused by Brsk2 elevation. (A and B) Insulin tolerance tests (ITTs, A) and blood glucose (B) were examined in Tg and WT mice after 4d, 2w, and 4w Dox-administration. (C to E) Body weights (C), gain of weights (D) and random blood glucose levels (E) of βKO and Ctrl mice fed with HFD. (F to H) ITTs (F), IPGTTs (G) and OGTTs (H) assays were performed in βKO and Ctrl mice fed with HFD for 12 weeks. (I to P) Hyperinsulinemic-euglycemic clamps in Tg and WT mice with Dox-induction for 1 week. Schematic representation of the experimental procedure (I). Steady GIR (J), whole glycolysis (K), glucose disposal rate (GDR, L), HGP (M), hepatic glucose uptake (N), synthesis of hepatic glycogen (O), and whole lipogenesis/glycogenesis (P) during hyperinsulinemia-euglycemic clamps. (Q) Pyruvate tolerance tests (PTTs) were performed in Tg and WT mice with Dox induction for 1 week. (R to T) PTTs (R) and phosphor-AKT(S308) levels (S) in βKO mice and Ctrls with HFD feeding for 12 weeks. Grey intensity of pAKT/AKT in Supporting Information Fig S (T). Data were presented by Mean ± SEM. n = 3-8 per group. n.s. = not significant, \**P <* 0.05, \*\**P <* 0.01.

However, the Tg Dox+ mice barely suffered from hyperinsulinemia-caused obesity. We therefore introduced HFD model to interrogate the contribution of Brsk2 to obesity and insulin resistance by using βKO mice. The βKO mice gained less body weights than Ctrl during HFD feeding (Fig 3 C and D). Random blood glucose levels were lower in the βKO mice (Fig 3E). Insulin sensitivity and glucose tolerance were greatly improved in the βKO mice (Fig 3 F-H). βKO mice also tolerated long-term HFD feeding (six months) (Supporting Information Fig S3C) as well as aging-induced hyperinsulinemia, overweight, glucose intolerance, and GSIS defects (Supporting Information Fig S3D). Thus, deletion of Brsk2 in mature β cells protects mice from obesity, T2DM as well as age-associated diabetes.

To find out the specific insulin resistance tissues caused by Brsk2 elevation, a hyperinsulinemia-euglycemic clamp was performed (Fig 3I). After constant insulin infusion, euglycemia was maintained by adjusting the GIRs, which significantly reduced in Tg Dox+ mice (Fig 3J and Supporting Information Fig S3E), indicating severely insulin resistant. Whole glucose metabolism and disposal rate (GDR) were reduced in the Tg Dox+ mice (Fig 3, K and L). Hepatic glucose production (HGP) was barely inhibited by hyperinsulinemia in Tg Dox+ mice (Fig 3M), consistent with refeeding hyperglycemia. The increased HGP was largely associated with an increase in hepatic glucose uptake and glycogen turnover (Fig 3, N and O and Supporting Information Fig S3F). While their whole-body lipogenesis was diminished despite increased glucose uptake in adipose tissues (Fig 3P and Supporting Information Fig S3G). Indeed, genes involved in de novo lipogenesis were significantly decreased in both liver and adipose tissue of Tg Dox+ mice (Supporting Information Fig S3H). This lipogenesis defect finally caused adipose atrophy of Tg Dox+ mice (Supporting Information Fig S3I). Thus, the hyperinsulinemia-euglycemic clamp results indicate that increased Brsk2 expression in β cells promotes glucose storage as hepatic glycogen but not conversion into fat, thereby giving rise to an increase in hepatic glucose overproduction and hyperglycemia in Tg Dox+ mice.

Net hepatic glucose production by pyruvate tolerance tests (PTTs) showed that Tg Dox+ mice produced more glucose than WT Dox+ mice (Fig 3Q). On the contrary, deletion of Brsk2 significantly inhibited HGP in HFD-feeding mice (Fig 3R). The improvements of tissue insulin sensitivities were also observed by increased pAKT levels in liver and WAT, but not in skeletal muscle (Fig 3 S and T). The gene expression levels revealed substantially improved de novo lipogenesis and reduced inflammation in βKO livers (Supporting Information Fig S3I). While no alterations in gluconeogenesis genes were detected (Supporting Information Fig S3I).

Taken together, these data involving acute and chronic induction or ablation of Brsk2 in mature β cells strongly support the notion that β cells can dominate systematic glucolipid metabolism by regulating Brsk2 expression. Mice with elevated Brsk2 mimic some of the defects caused by HFD feeding in terms of insulin resistance manifested as hepatic glucose overproduction and adipose dysfunction.

### Parasympathetic input enables β-cell oversecretion via Chrm3/Brsk2/Plaa axis

The underlying molecular mechanism resulting in hyperinsulinemia before the onset of frank T2DM was investigated by RNA sequencing of mouse islets. In total, of 23,318 gene transcripts were captured, and 1,020 genes showed significant expression differences (p-value < 0.05 and > 2-fold change) between the two groups (Fig 4A). KEGG enrichments of upregulated gene sets were linked to pathways associated with Pancreatic secretion, Neuroactive ligand-receptor interaction, and among others (Fig 4B and Table S1), indicating neuroregulation of insulin secretion.

**Figure 4.**
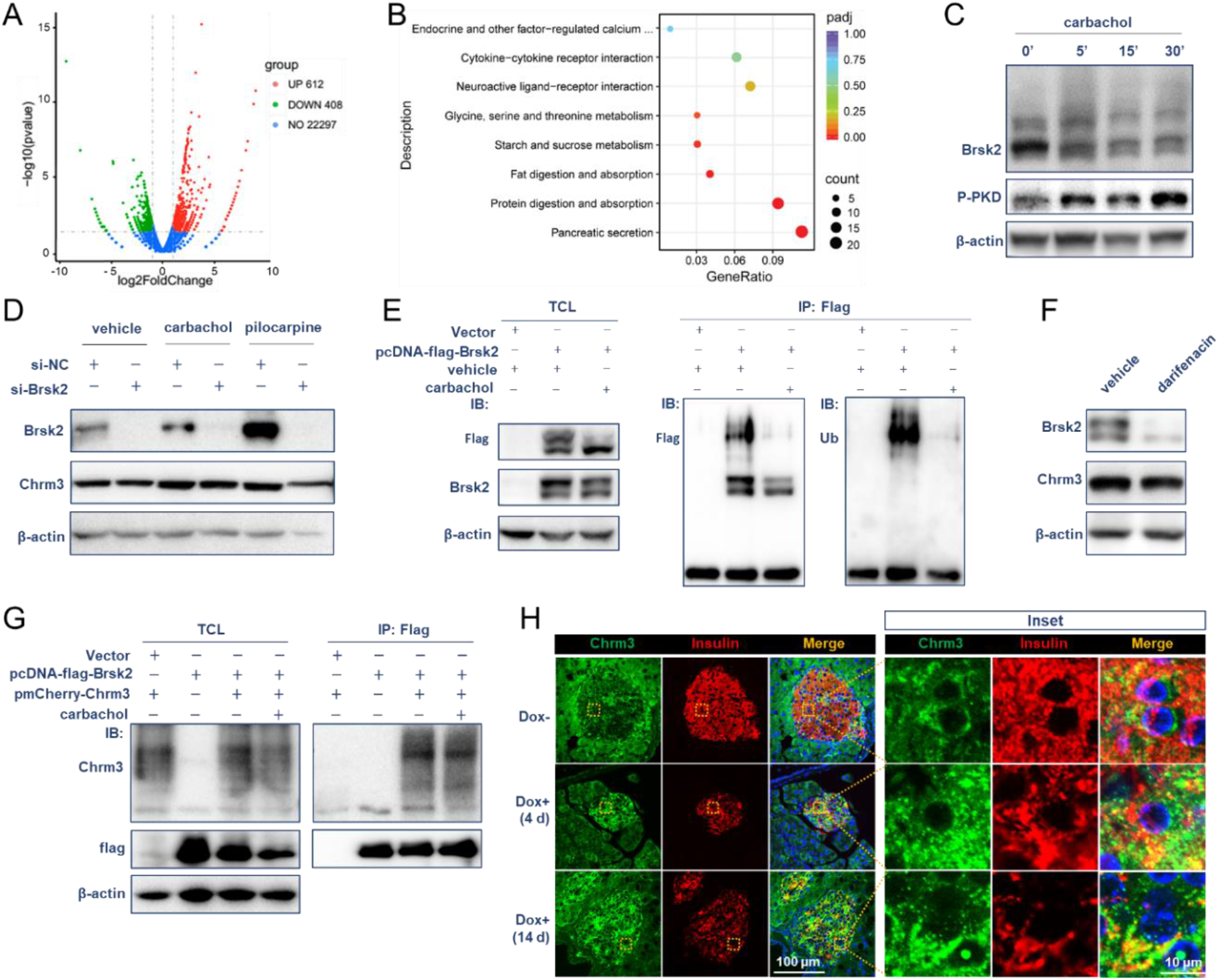
Parasympathetic input stabilizes Brsk2 through Chrm3. (A and B) Primary islets from WT and Tg mice with Dox induction for 4 days were isolated for RNA-sequencing. Volcano plots and top eight KEGG enrichments of upregulated genes were displayed. (C) Immunoblots show protein levels of Brsk2 and P-PKD in MIN6 cells treated with 10 μmol/l carbachol. (D) MIN6 cells were transected with si-*Brsk2* or si-NC for 72 h and treated with carbachol (10 μmol/l), pilocarpine (100 μmol/l) for additional 1 h for protein detection. (E) MIN6 cells were transfected with pcDNA-flag-Brsk2 for 24 h and treated with carbachol (10 μmol/l) for 30 min. Cells were collected for co-immunoprecipitation (co-IP) assays by using anti-flag antibodies, then protein samples of total cell lysate (TCL) and co-immunoprecipitates were analyzed. (F) MIN6 cells were treated with or without darifenacin (10 μmol/l) for 1 h, and protein levels of Brsk2 and Chrm3 were detected. (G) HEK293T cells were co-transfected with pcDNA-flag-Brsk2 and pmCherry-Chrm3 for 24 h, and treated with carbachol (10 μmol/l) for additional 30 min. Co-IP assays were performed by using anti-flag antibodies, and protein levels of flag-Brsk2, mcherry-Chrm3 in TCL and co-immunoprecipitates were analyzed. (H) Pancreatic slices of Tg Dox+ mice were co-stained with Chrm3 (green), insulin (red) and nuclei (blue). Representative images were shown. Right panels were enlarged pictures. Data were presented by Mean ± SEM. n = 3-16 per group. n.s. = not significant, \**P <* 0.05, \*\**P <* 0.01.

We first tested the contribution of sensory neurons since olfactory transduction ranked first among the top altered pathways (Supporting Information Fig S4 A and B). Rozita et al. reported that capsaicin-sensitive Trpv1^+^ sensory neurons contributed to β-cell function and diabetes etiology (Razavi, Chan et al., 2006). However, capsaicin pretreatment was unable to prevent Tg Dox+ mice from hyperglycemia (Supporting Information Fig S4 C and D), while gene expression levels of olfactory receptors (ORs), such as Olfr78 and Olfr1420 were significantly increased in Tg Dox+ islets, as well as in HFD islets, which could be canceled by Brsk2 deletion (Supporting Information Fig S4 E and F). Previous reports have shown that sympathetic outflows could inhibit insulin secretion, whereas parasympathetic outflows could increase insulin release (Rodriguez-Diaz & Caicedo, 2014). Indeed, parasympathetic acetylcholine analog, carbachol significantly upregulated OR gene expressions, while Brsk2 knockdown inhibited the carbachol-potentiated OR levels (Supporting Information Fig S4 G and H), suggesting that olfactory transduction was dependent on a parasympathetic acetylcholine/Brsk2 pathway.

Next, a direct response of Brsk2 expression to parasympathetic outflows was determined. Carbachol treatment enhanced the phospho-PKD level (Fig 4B), consistent with previous studies (Sumara, Formentini et al., 2009). Carbachol treatment also intensified the smearing of Brsk2 band (Fig 4B). Beta cells have at least two muscarinic cholinergic receptors, Chrm3 and Chrm4, which trigger insulin release via Chrm3-Gq-Ca^2+^ pathway and Chrm4-Gs-cAMP pathway, respectively. Only Chrm3 agonist pilocarpine treatment triggered Brsk2 band smearing (Supporting Information Fig S4I). Moreover, treatment with carbachol relocalized EGFP-Brsk2 complex from the plasma membrane and cytosol to the lateral membrane where insulin is released (Supporting Information Fig S4 J and K).

Alternatively, prolonged carbachol or pilocarpine treatments increased protein level of Brsk2 due to removal of polyubiquitination modification of Brsk2 (Fig 4, D and E). Conversely, darifenacin, a specific Chrm3 antagonist, reduced Brsk2 protein amount via enhancing K48-linked polyubiquitination (Fig 4F and Supporting Information Fig S4L). The reciprocated regulation between Chrm3 and Brsk2 was also observed (Fig 4D), probably because Brsk2 was interacted with Chrm3 (Fig 4G and Supporting Information Fig S4M). In Tg Dox+ mice, Chrm3 was enhanced by acute Brsk2 overexpression, distributed from perinuclear region to cytoplasm in dot shape and located at lateral membrane (Fig 4H), akin to the cellular distribution of Brsk2 post-carbachol treatment. Meanwhile, insulin was co-stained with Chrm3 and significantly decreased by Brsk2-elevation (Fig 4H). Combined with serum high levels of C-peptide and proinsulin, we speculated that parasympathetic acetylcholine stabilized Brsk2 which triggered persistent exocytosis in a Chrm3-dependent pathway.

Oral feeding triggers autonomic outflows and regulates the cephalic phase of insulin secretion before nutrient absorption, while i.p. glucose injection bypasses any autonomic effects. The in vivo effect of Chrm3 and Brsk2 interaction was therefore investigated by oral GSIS assays. Despite Tg Dox+ mice had lower insulin content in pancreas (Fig 5A), oral glucose simulated more insulin at cephalic phase, while darifenacin blocked this insulinotropic effect at both cephalic and glucose absorptive phases (Fig 5B). HFD mice with hyperinsulinemia responded more strongly to carbachol-triggered insulin secretion than NCD mice as well as HFD mice with euinsulinemia, independent of glucose absorption (Supporting Information Fig S5 A and B), akin to Brsk2-elevated mice. Interestingly, β-cell hypersecretion in HFD mice significantly inhibited by pretreatment of three autonomic inhibitors (Supporting Information Fig S5C). Deletion of Brsk2 blocked insulin release at cephalic phase but not at the glucose absorption phase in both NCD-fed and HFD-fed conditions (Fig 5C). Thus, the in vivo evidence suggests that oral feeding evokes cephalic insulin secretion via acetylcholine/Chrm3/Brsk2 pathway, thereby contributing to the hyperinsulinemia under HFD condition.

**Figure 5.**
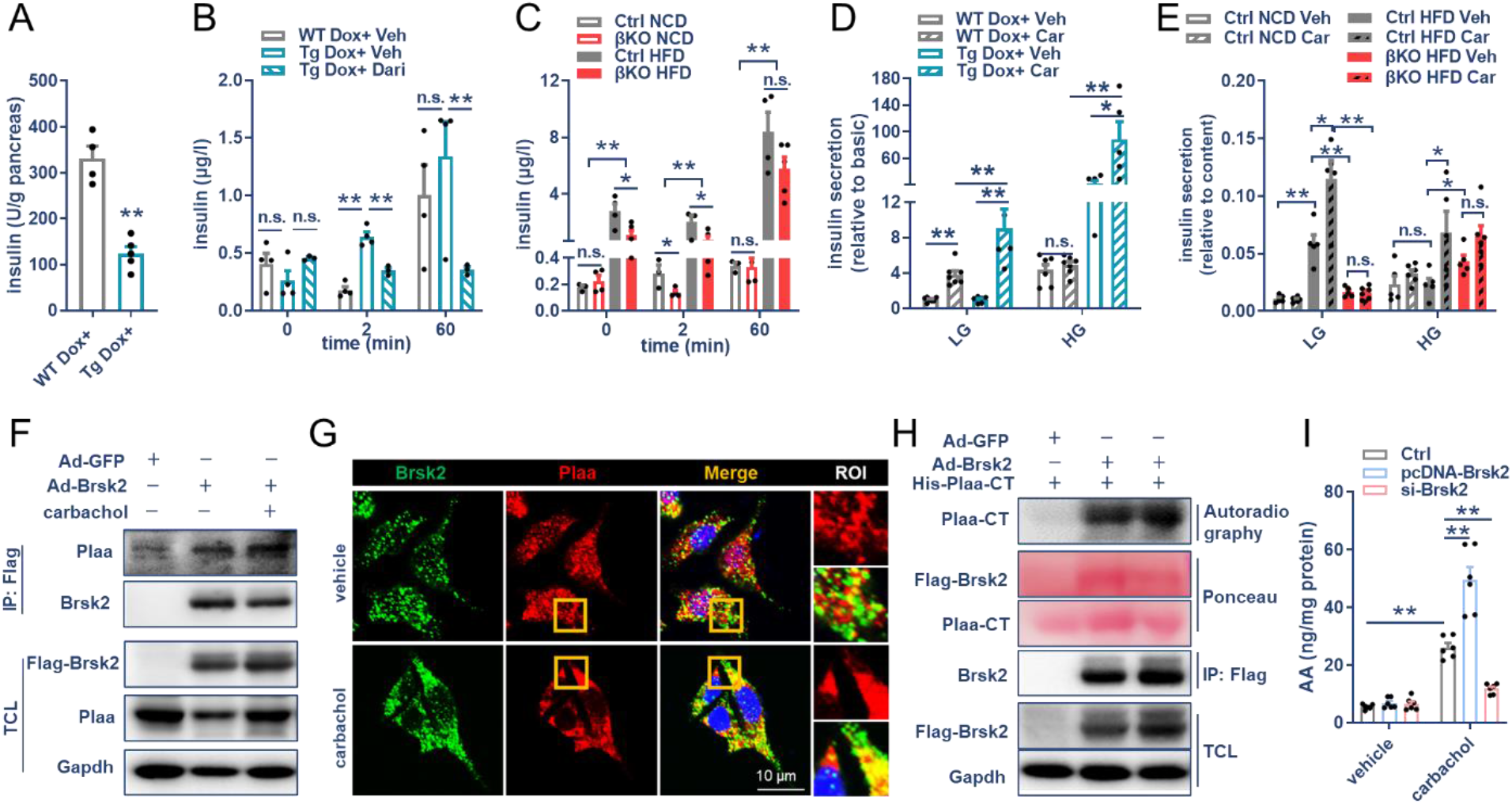
Brsk2 promotes β-cell oversecretion via Chrm3/Brsk2/Plaa axis. (A) Insulin content of whole pancreas extracts from WT and Tg mice after 4-week Dox-induction. (B) WT and Tg mice with 4-week Dox-induction were ip. injected with darifenacin (10 mg/kg) or vehicle for 30 min and then administered with glucose (2mg/ml) intragastrically. Serum insulin levels were detected. (C) Ctrl and βKO mice with HFD for 12 weeks and subjected to OGTTs for serum insulin measurement. (D and E) Primary islets from WT and Tg mice with Dox-induction for 4 weeks (D) and from Ctrl and βKO mice with HFD for 12 weeks (E) were performed GSIS assay with or without carbachol stimulation. (F) MIN6 cells were infected with Ad-Brsk2 for 24 h and treated with carbachol (10 μmol/l) for 30 min. Cells were collected for co-IP assays by using anti-flag antibodies. (G) MIN6 cells were treated with carbachol for 30 min and then co-stained with Brsk2 (green), Plaa (red) and nuclei (blue). ROI: region of interest. (H) HEK293T cells were infected with Ad-Brsk2 for 24 h. Brsk2 protein was enriched by using anti-flag antibodies. Flag-Brsk2 was used for protein kinase assay by using 6*His-Plaa-CT (human Plaa, 496-795 amino acids) as a substrate. Kinase and substrate reacted samples were prepared and Plaa phosphorylated by Brsk2 was detected by autoradiography. Ponceau S staining detected the protein levels of Plaa-CT and Flag-Brsk2 in kinase reaction. (I) MIN6 cells were transfected with pcDNA-Brsk2, si-Brsk2 or relative control (Ctrl) for 48 h and treated with carbachol for 1 h. Cells were collected for detection of arachidonic acid (AA) by ELISA. β-actin or Gapdh was used as an internal standard for immunoblots. Data were presented by Mean ± SEM. n = 3-16 per group. n.s. = not significant, \**P <* 0.05, \*\**P <* 0.01.

The complexity of in vivo results was circumvented by isolating primary islets for ex vivo assays and introducing carbachol to mimic in vivo parasympathetic activation. Carbachol treatment released more insulin by Tg Dox+ islets and 4-month HFD islets, which remained much less insulin content (Fig 5D and Supporting Information Fig S5 D and E). While, carbachol did not affect the insulin content in the WT Dox+ islets or in 2-month HFD islets (Supporting Information Fig S5 D and E). A strengthened insulin secretion by carbachol treatment was also avoided by Brsk2 deletion (Fig 5E, and Supporting Information Fig S5 F-H). Taken together, our in vivo and in vitro data demonstrated that a parasympathetic acetylcholine/Chrm3/Brsk2 axis links oral feeding to enhanced islet olfactory transduction, cephalic insulin release, and reductions in pancreatic insulin content.

To find out the downstream substrates of acetylcholine/Chrm3/Brsk2 axis, Brsk2 co-immunoprecipitated (co-IP) proteins with or without carbachol treatment were identified by mass-spectrometry (Supporting Information Fig S5 I to K and Table S2). KEGG pathways enriched Protein processing in endoplasmic reticulum, insulin resistance, and type 2 diabetes ((DAVID Bioinformatics Resources 6.8, Supporting Information Fig S5L). One candidate was phospholipase A2 activating protein (Plaa) which can assist phospholipase A2 to generate arachidonic acid (AA), an insulinotropic polyunsaturated fatty acid (Tsunoda & Owyang, 1994). Interaction between Brsk2 and Plaa was confirmed (Fig 5 F and G). Consistent with Brsk2, Plaa amount also increased in HFD mouse islets (Supporting Information Fig S5M). Interference of Plaa significantly decreased insulin secretion under both glucose-and carbachol-stimulated conditions (Supporting Information Fig S5 N to P). Notably, Plaa was phosphorylated by Brsk2 and facilitated carbachol-induced AA generation (Fig 5 H and I and Supporting Information Fig S5Q). Above results indicate that parasympathetic outflows stimulate AA production to cause β-cell oversecretion via Chrm3/Brsk2/Plaa axis.

### Blocking the parasympathetic input to pancreas remits diabetes via inhibiting HGP

The link between oral feeding and islet PNS signal transduction amplified by Brsk2 overexpression was severed by restoring the Brsk2 level via Dox withdrawal, transplantation of denervated islets, autonomic antagonist administration, and vagotomy surgery.

The levels of random blood glucose in Tg Dox+ mice started to increase at day 4 post Dox-induction and reached to around 400 mg/dL at day 9, then maintained this peak during the observation (Fig 6A). Surprisingly, Tg Dox+ mice showed normoglycemic levels immediately after Dox withdrawal, regardless of whether Dox was removed at an early (days 9 and 37) or late (day 147) stage (Fig 6A). Dox withdrawal inhibited OGTT-caused cephalic insulin secretion and recovered IPGTT insulin secretion (Fig 6B), thereby largely conserving whole pancreas insulin amounts (Fig 6C). Insulin staining recovered from the uneven pattern and became uniformly distributed as the Brsk2 level restored (Fig 6D and Supporting Information Fig S6A).

**Figure 6.**
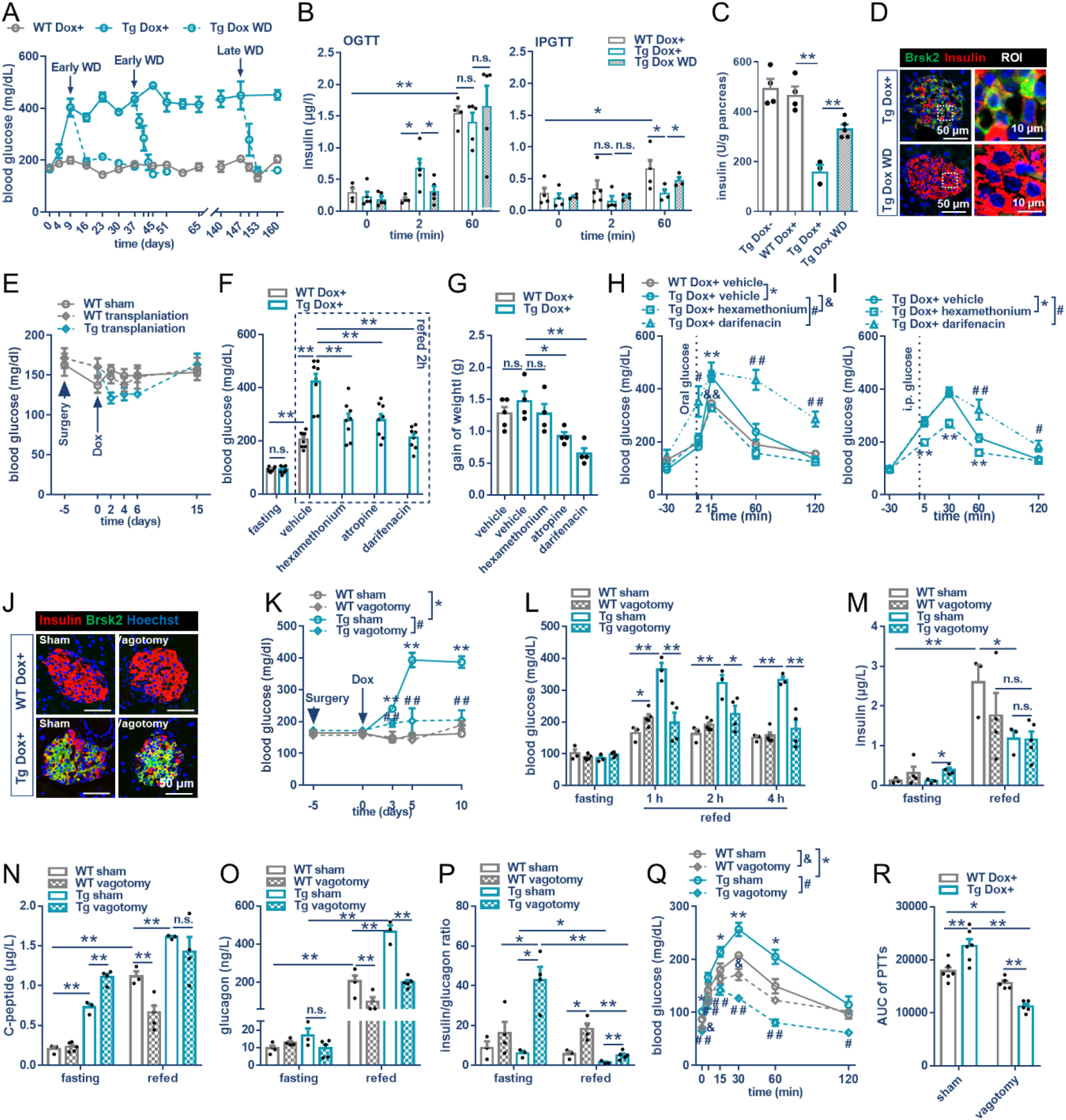
Blocking the parasympathetic input to pancreas remits diabetes via inhibiting HGP. (A) WT Dox+ and Tg Dox+ mice were drinking normal water at day 9 and 37 (early withdraw, early WD), or day 147 (late WD). Random blood glucose levels were monitored. (B to D) Mice with 4-week Dox-induction and 4-week Dox-WD were subjected to OGTTs and IPGTTs for serum insulin detection (B), pancreas insulin content (C), and immunostaining for insulin (green), G/S/PP (red), and nuclei (blue) (D). (E and F) Fourteen-hour fasted mice were pretreated with indicated drugs for 30 min and refed for 2 h, and blood glucose level and gain of weights were analyzed. (G and H) Tg Dox+ or WT Dox+ mice, pretreated with indicated drugs for 30 min were subjected to oral-(G) or i.p. injected (H) GTTs. (I) Mouse islets from WT and Tg mice were transplanted into subcapsular kidney of WT mice. Recipient mice were administrated with Dox and random blood glucose levels were examined. (J to R) Tg and WT mice were subjected to vagotomy or sham surgery and then administrated with Dox-containing water. Immunostaining of Insulin (red), Brsk2 (green), and nuclei (blue) was performed in pancreas slices post 21-day Dox induction (J). Random blood glucose levels were monitored (K). (L to P) Serum levels of fasting and refed glucose (L), insulin (M), C-peptide (N), glucagon (O), and ratio of insulin to glucagon (P) in fasting and 2-h refed vagotomy mice after Dox-administration for 6 days. PTTs of vagotomy mice after Dox-administration for 11 days (Q) and AUC of Q (R). Data were presented by Mean ± SEM. n = 4-8 per group. n.s. = not significant, *^, #^ or ^&^ *P <* 0.05, **, ^##^ or ^&&^ *P <* 0.01.

Parasympathetic, sympathetic, and olfactory nerves that reach to islets are truncated during islet transplantation, after which it takes more than one month for nerve regeneration. We took advantage of this specific time window to study whether islet innervations contributed to T2DM caused by Brsk2 overexpression. Islets isolated from Tg and WT mice were transplanted into the subcapsular kidneys of WT mice. The successful Brsk2 induction by Dox-water in the kidney-transplanted Tg islets was confirmed by H&E and Brsk2 staining (Supporting Information Fig S6B). Compared to the sham group, the body weights, blood glucose levels of random and fasting of islet transplanted mice barely changed, while better postprandial glucose control due to the additional insulin secreted by the transplanted islets (Fig 6E and Supporting Information Fig S6, C to E). However, no hyperglycemia was observed in the Tg islet transplanted mice.

Denervation causes mixed effects on the autonomic efferents. We examined which autonomic maladjustments influenced the hyperglycemia phenotype caused by Brsk2 elevation. Autonomic blockages greatly normalized the blood glucose levels upon refeeding, with hexamethonium showing the greatest improvement, independent of weight gain (Fig 6 F and G). Pretreatment with hexamethonium enhanced the glucose-clearance efficiencies (Fig 6 H and I), and this was largely attributed to an early inhibition of glucagon secretion and barely altered insulin secretion despite the late inhibition of β-cell secretions (Supporting Information Fig S6 F and G). By contrast, darifenacin pretreatment exaggerated the glucose excursions in the Tg Dox+ mice (Fig 6 H and I), probably due to the continuous inhibition of β-cell secretions regardless of simultaneous inhibition of α-cell secretions (Supporting Information Fig S6 H and I). These results further support the conclusion that parasympathetic outflows trigger islet hormonal oversecretions, and that inhibition of glucagon, but not insulin effects could abate the refeeding hyperglycemia.

Nonspecific effects of the autonomic antagonists were excluded by subdiaphragmatic vagotomy. Gastric distension was considered a positive sign of a successful surgery (Supporting Information Fig S6 J and K). Immunostaining confirmed that vagotomy did not affect the Dox induction of Brsk2 expression in β cells (Fig 6J). Vagotomy prevented random and refeeding hyperglycemia in the Brsk2-overexpressing mice (Fig 6 K and L). In fact, vagotomized mice showed increased body weights and decreased blood glucose levels due to increased fasting insulin levels (Supporting Information Fig S6 L and M). The euglycemic effect of vagotomy in Brsk2-overexpressing mice was attributed to increased fasting insulin and C-peptide levels, combined with decreased glucagon levels after refeeding (Fig 6 M to O).

Careful calculations of the insulin/glucagon ratio at single mouse level revealed that vagotomy increased this ratio in the WT Dox+ mice after refeeding, whereas vagotomy increased insulin/glucagon ratio in both the fasting and refed conditions in the Tg Dox+ mice (Fig 6P), thereby improving the HGP in the vagotomy groups (Fig 6 Q and R). Hepatic gluconeogenesis was excluded while energy storage as fat was implied to the metabolic improvements of vagotomy (Supporting Information Fig S6 N and O). Therefore, vagotomy surgery, which directly blocks the parasympathetic innervation to islets, also prevents Tg Dox+ mice from developing T2DM due to inhibition of hepatic glucose overproduction and adipose atrophy by optimizing the insulin/glucagon ratios.

Taken together, the data presented here demonstrate that Brsk2 overexpression amplifies parasympathetic signal transduction in β cells, which in turn causes β-cell exocytosis and reduces pancreas insulin content, which is likely associated with the increased serum glucagon level postprandially. Blocking the link between the parasympathetic outflows and the islets can therefore remit T2DM.

### Inhibition of Brsk2 treats HFD-induced mouse T2DM

CGP 57380 is a verified inhibitor of Brsk2 (Grzmil, Huber et al., 2014). Treatment with CGP 57380 significantly suppressed Brsk2 protein levels in MIN6 cells (Fig 7A), while had no effect on MIN6 cell viability (Supporting Information Fig S7A). Since no alteration of phosphor-AMPK was observed, the decreased P-AMPK substrate phosphorylation levels were attributed to the reduced Brsk2 levels (Fig 7A and Supporting Information Fig S7B). Inhibition of Brsk2 enhanced GSIS function in MIN6 cells (Fig 7B). Human islet perfusion indicated that the enhanced insulin secretion was attributed to enlarged immediate releasing pool of insulin granules (Fig 7 C and D).

**Figure 7.**
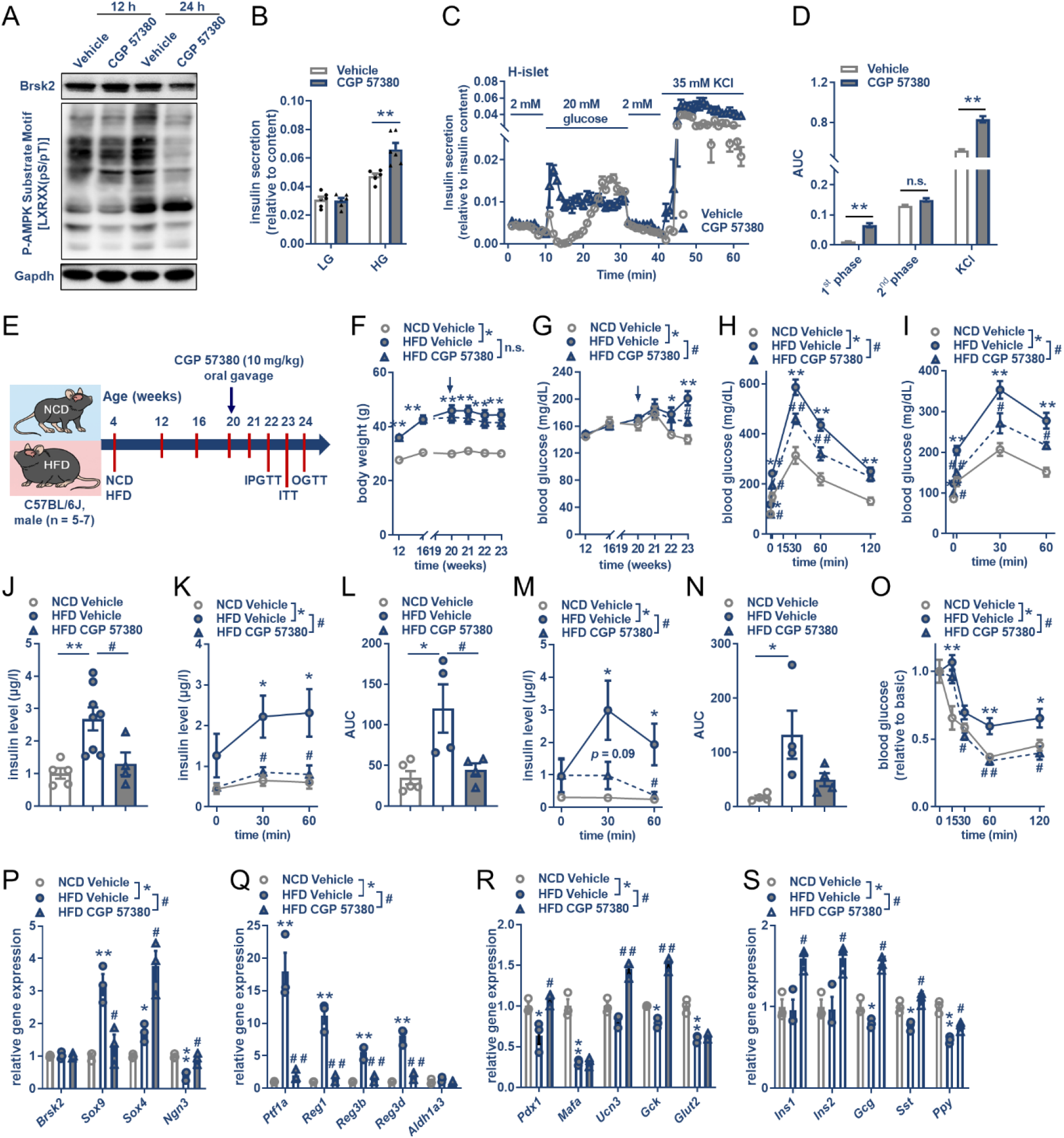
Inhibition of Brsk2 treats HFD-induced mouse T2DM. (A) MIN6 cells were treated with CGP 57380 (10 μM) for indicated time and protein levels of Brsk2 and P-AMPK Substrate Motif were detected. (B) MIN6 cells were treated with CGP 57380 (10 μM) for 24 h and GSIS assays were performed. (C) Primary human islets were treated with CGP 57380 (10 μM) for 24 h and islet perfusion assays were performed. (D) AUC of GSIS at first phase and second phase, and K+-stimulated insulin release. (E to S) C57Bl/6J mice aged 4 weeks were subjected to NCD or HFD for 16 weeks, by which time HFD mice were administrated with CGP 57380 (10 mg/kg) through oral gavage once every day. Schematic of CGP 57380 treatment on HFD mice (E). Body weights (F) and RBG levels (G) were monitored. IPGTTs (H), OGTTs (I), random serum insulin levels (J), insulin secretion during IPGTTs (glucose: 2g/kg, K) and OGTTs (glucose: 2g/kg, M), and ITTs (O) assays were performed. AUC of K (L) and AUC of M (N) were calculated. (P to S) Gene expressions of islets isolated from NCD with vehicle, and HFD with or without CGP 57380 treatment for one month. Data were presented by Mean ± SEM. n = 3-8 per group. n.s. = not significant, \**P <* 0.05, \*\**P <* 0.01.

We took advantage of this drug to test the anti-diabetic effect on 16-week HFD-induced mice. Mice were subjected to CGP 57380 (10mg/kg) administration once every day and metabolic characteristics were monitored (Fig 7E). CGP 57380 administration had no effect on body weight and minimal effects on RBG levels (Fig 7 F and G). However, CGP 57380 significantly inhibited the fasting blood glucose levels and largely improved glucose disposal abilities on HFD feeding mice, due to the improvement of insulin sensitivity (Fig 7 H to J). CGP 57380 also restricted the increased insulin secretion demand caused by HFD via improving glucose sensing abilities (Fig 7 K to O).

To define the mechanisms of metabolic improvements from Brsk2 inhibition, RNA-sequencings of primary islets isolated from Vehicle or CGP 57380 administrated HFD mice were introduced. Overall, 1431 genes were significantly altered with p-value < 0.05 and >2-fold change (Supporting Information Fig S7C). GSEA analysis revealed that top 5 KEGG pathways were Notch signaling pathway, Pancreas secretion, Fat and protein digestion and absorption, and Endocrine calcium absorption (Supporting Information Fig S7E). Notably, Olfactory transduction and Arachidonic acid metabolism were also enriched (Supporting Information Fig S7 D and E), implicating that HFD-induced metabolic abnormities were akin to Brsk2 elevation and those defects could be eliminated by pharmacological inhibition of Brsk2. We also confirmed some sets of genes by qRT-PCR assays. Gene expression levels in islets showed HFD-feeding increased pancreas progenitor genes, exocrine genes, while decreased endocrine genes and hormone genes, all of which could be rescued by CGP 57380 treatment (Fig 7 P to S). Thus, pharmacological inhibition of Brsk2 can treat HFD-induced T2DM and ameliorate β-cell failure.

### Human BRSK2 loci mutations are associated with worsening glucose metabolism

In this part, we further analyzed the human gene BRSK2 with metabolic traits in Shanghai Nicheng Cohort Study (Table S3). After screening genetic variants in BRSK2, we identified 3 variants associated with diabetes mellitus and related glucose metabolic traits, including rs112377266 (->dup(G)_4_CTCACCTGTGG, an insert variant in intro 15 of BRSK2 gene, minor allele frequency = 0.37), rs61002819 (G>-, a deletion variant in splice region, variant frequency = 0.11), a deletion variant in splice region, variant frequency = 0.11) and rs536028004 (C>T, a 3 prime UTR variant, variant frequency = 0.01). The linkage disequilibrium relationship was shown in Table S4.

Of note, subjects with minor allele have higher risk of T2DM, indicating the important role of BRSK2 in glucose metabolic process. As shown in Fig 8A and Supporting Information Fig S8A, Dup(G)_4_CTCACCTGTGG allele of rs112377266 carries had higher incidence of T2DM (-/- homozygote: 34.55%, -/dup(G)_4_CTCACCTGTGG heterozygote: 36.78%, dup(G)_4_CTCACCTGTGG homozygote: 42.53%). We also analyzed the plasma glucose and insulin level after 75g OGTT at 0, 30 and 120 min, the results showed that dup(G)_4_CTCACCTGTGG carriers had higher plasma glucose and plasma insulin, indicating worse glucose metabolism. Logistic regression and multiple linear regress showed that rs112377266 was associated with T2DM (OR&95%CI = 1.04 [1.02-1.05], P = 2.42×10^−5^) and Stumvoll 2^nd^ of insulin secretion (β = -3.19, SE = 0.97, p = 9.87×10^−4^) after adjusting for age and sex. Fig 8B and Supporting Information Fig S8B showed that another splice region variant rs61002819 also associated with T2DM (OR&95%CI = 1.13 [1.04-1.05], P = 9.84×10^−4^), subjects with deletion had higher incidence of T2DM (G/G homozygote: 36.07%, G/- heterozygote: 38.64%, -/- homozygote: 47.37%). GUTT-ISI index showed insulin insensitivity in G>-deletion carriers. Furthermore, Fig 8C and Supporting Information Fig S8C showed that rs536028004 located in 3 prime UTR regions was a relatively low frequency variant, T allele carriers had higher incidence of T2DM (CC homozygote: 36.38%, T allele carriers: 49.07%), glucose intolerance and insulin resistance. After adjustment for age and sex, rs536028004 was also associated with T2DM (OR&95%CI = 1.03 [1.01-1.06], P = 9.90×10^−3^).

**Figure 8.**
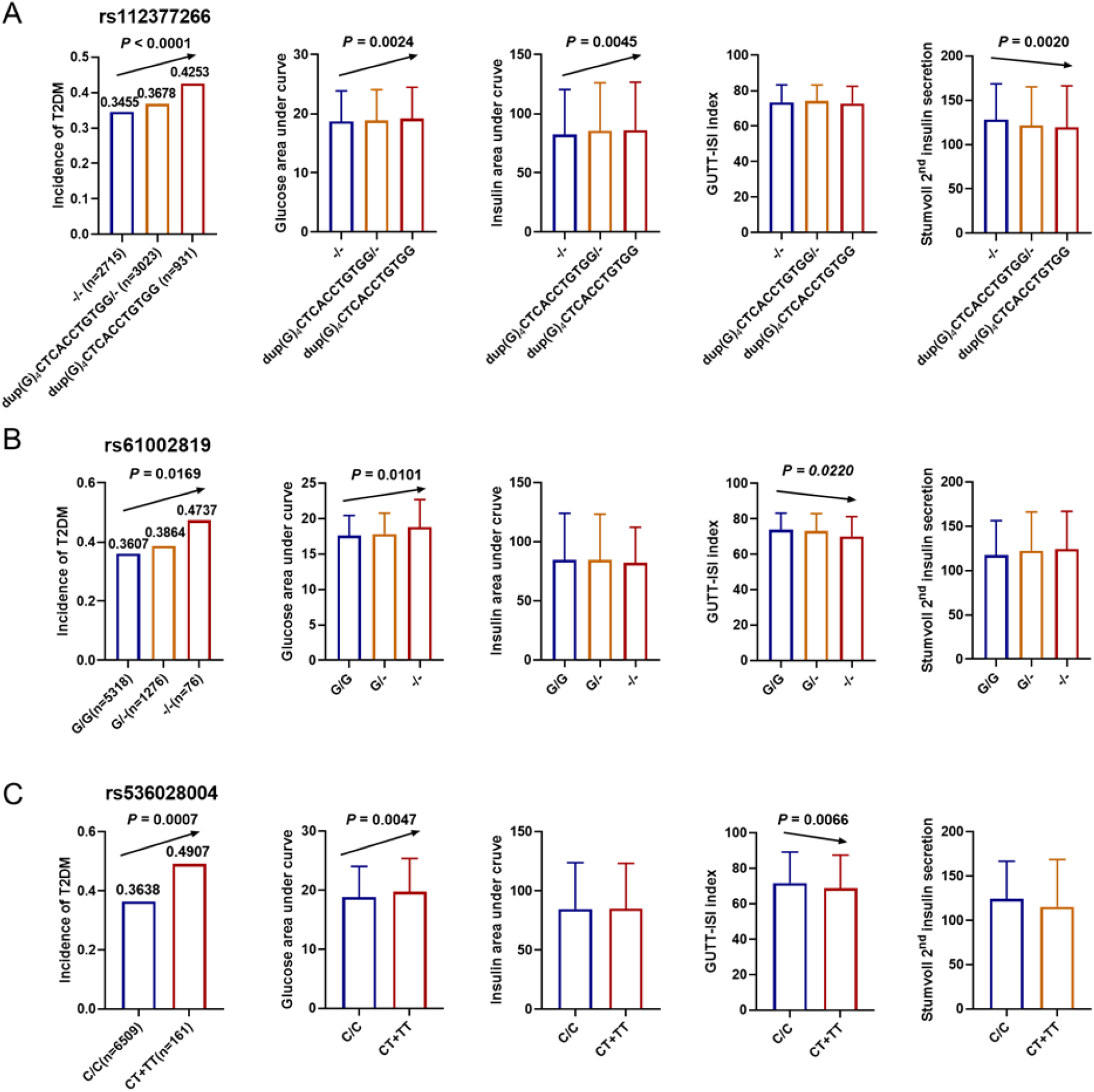
Human *BRSK2* loci mutations are associated with worsening glucose metabolism. (*A, B* and *C*) Association between *BRSK2* loci genotyping (rs112377266, rs61002819 and rs536028004) and glucose metabolism. Each panel (from left to right) showed the incidence of T2DM, the area under curve of plasma glucose level after OGTT, the area under curve of plasma insulin level after OGTT, GUTT-ISI index and Stumvoll 2^nd^ insulin secretion index.

Above results reveal that BRSK2 mutations have potential roles on regulating glucose metabolism in human.

## DISCUSSION

Insulin is the only hormone that lowers blood glucose levels, and its synthesis and secretion by β cells is harmonized by circulatory and neural signals perceived by the islets in the context of intermittent feeding. Alterations in diet composition, such as an HFD, can promote inappropriate insulin secretion by unrecognized reasons. Here, we found that Brsk2 can sense parasympathetic signals to initiate preabsorptive or cephalic insulin release. If amplified by Brsk2 elevation, such as in HFD islets, these signals can induce persistent β-cell exocytosis, thereby causing hyperinsulinemia, obesity, insulin resistance and T2DM via Chrm3-Brsk2-Plaa axis (Figure 9, Graphic abstract).

**Figure 9.**
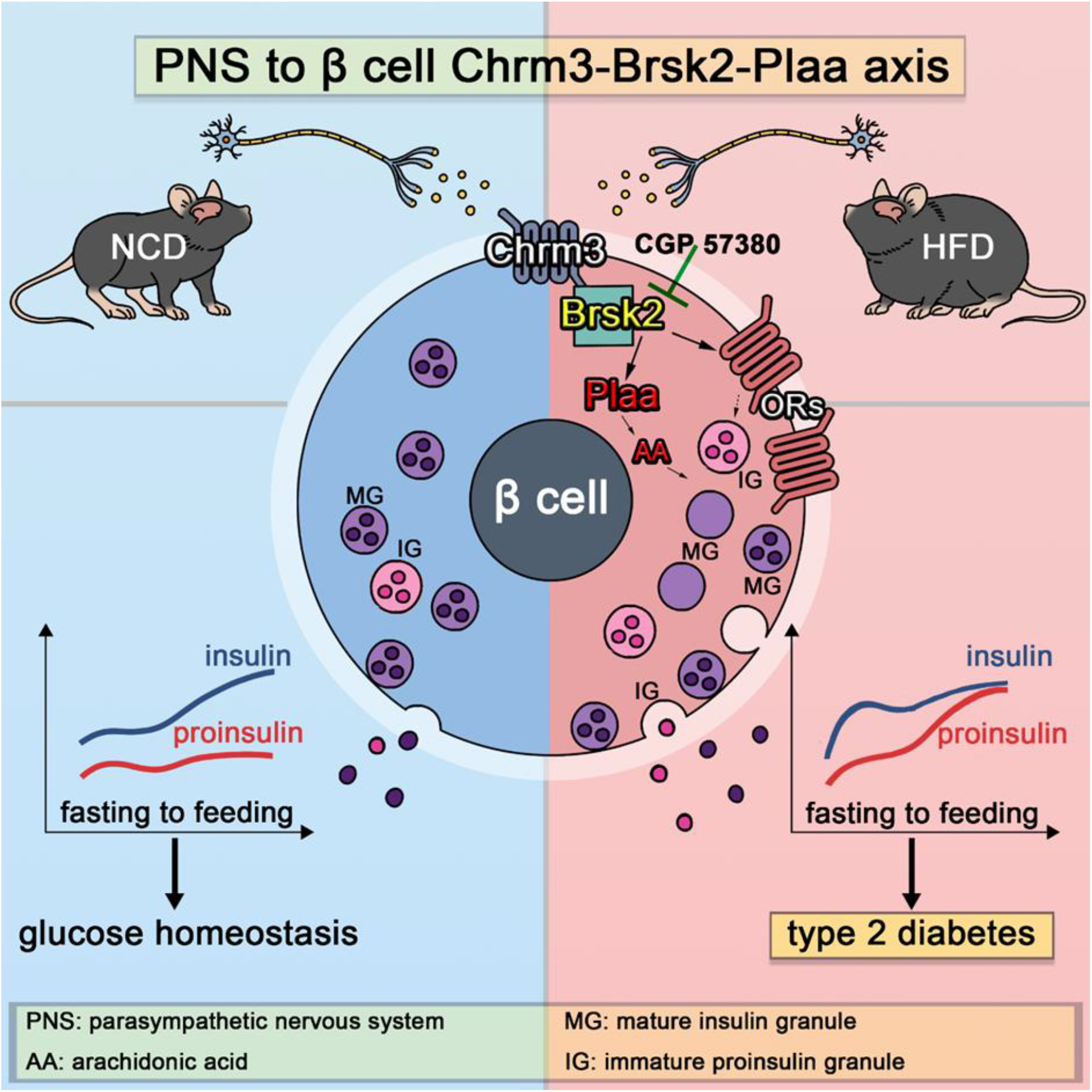
Graphical abstract.

Previous studies have shown that mice with whole body or pancreatic knockout of Brks2 were deficient in islet mass (Nie et al., 2013c), suggesting that Brsk2 is important for islet development. The neuro-insular network is imaged to coincide with stages of rapid growth, differentiation, and maturation in the embryonic pancreas (Burris & Hebrok, 2007). Due to its restricted tissue expression pattern in the brain and pancreas, the neuro-insular axis likely manages islet development, which is dependent on mutual Brsk2 expression. Genetic or pharmacological ablation of sympathetic innervation during mouse development results in altered islet architecture, reduced insulin secretion, and impaired glucose tolerance (Liggett, 2009). Parasympathetic innervation is also critical for establishment of the β-cell mass in the postnatal period and for long-term maintenance of β-cell function (Tarussio, Metref et al., 2014). Dysfunctions in either the sympathetic or parasympathetic branches compromise mature islet function. This therefore complicates the determination of whether the suppression of islet mass expansion following global or pancreatic deletion of Brsk2 originates from defects in one or in both pathways. Recently developed deep-tissue microscopy methods for 3-dimensional islet neurohistology may assist in resolving these issues (Alvarsson, Jimenez-Gonzalez et al., 2020). Nevertheless, β-cell Brsk2 may very likely respond to autonomic outflows at the embryonic stage.

In the present study, the islet mass in mice with the Brsk2 deletion in mature β cells was identical to that in their control littermates under both NCD and HFD conditions, extending our study in mature islets under metabolic stressed condition. Interestingly, Brsk2 expression was enhanced in mouse islets within one month of HFD feeding and was accompanied by overweight and hyperinsulinemia, while euglycemia. Brsk2 elevation was attributed to multiple phosphorylation of Brsk2 C-terminal modulated protein stability. Literatures have shown that PKC could be activated by Gq-coupled GPCRs including FFAs and acetylcholine (Inoguchi, Li et al., 2000, Oancea, Bezzerides et al., 2003). During HFD setting, mice exposed to high dose of FFAs and encountered to autonomic imbalanced conditions (Berthoud, Albaugh et al., 2021, O’Brien, Hinder et al., 2017). However, our previous study has shown that FFAs treatment only shifted Brsk2 band from 86 to 78kDa (Wang, Liu et al., 2019), whereas parasympathetic activation induced both band shift and an increase of protein level. Alternatively, HFD feeding could cause insulin resistance and AKT inactivation in mouse islets (Paschen, Moede et al., 2019). Here, AKT inhibitor also markedly increased Brsk2 protein level. Thus, mouse islets can directly respond to metabolic stress by increasing Brsk2 protein amount. Deletion of Brsk2 can protect mature mice from HFD-induced metabolic abnormities by lowering insulin secretion to maintain glucose-responsive insulin secretion. These results raise the likelihood of an association of diabetes predisposition with Brsk2 levels in adults. Indeed, acute induction of Brsk2 expression in mature β cells caused postprandial hyperinsulinemia and hyperglycemia within a week; while chronic Brsk2 overexpression triggered frank diabetes characterized by insulin resistance and β-cell dysfunction. Unexpectedly, regardless of whether Brsk2 was acutely or chronically induced, the mice recovered their normoglycemic state immediately after Brsk2 was returned to basic levels, suggesting that the amount of Brsk2 protein in β cells directly governs glucose homeostasis.

Comprehensive RNA-sequence profiling showed that enhanced olfactory transduction was largely associated with acute Brsk2 overexpression. Generally, olfactory inputs help to coordinate food appreciation and selection and they participate in neuronal circuit development within the central nervous system to integrate internal cues into autonomic responses via the sympathetic and parasympathetic efferents (Berger, Scheel et al., 2015, Brandt, Nolte et al., 2018, Meek, Nelson et al., 2016). Eliminating sensory fibers prevents insulitis and diabetes in type 1 diabetes-prone NOD mice (Razavi et al., 2006). However, eliminating TRPV1^+^ sensory neurons has minimal effects on Brsk2-overexpression induced T2DM. Further investigation reveals that the enhanced islet olfactory transduction by Brsk2 is attributed to parasympathetic activation. Since these alterations occur prior to the appearance of insulin resistance and hyperglycemia, increases in Brsk2 may indicate an attempt to remodel β-cell function, islet mass, and architecture as part of a response to mild metabolic stress. However, islet adaptation would entail hyperinsulinemia to promote insulin resistance and β-cell dysfunction and would inevitably lead to diabetes (Alarcon, Boland et al., 2016), as occurs in the Brsk2-overexpressing mice, probably due to parasympathetic triggered cephalic insulin oversecretion. Mice transplanted with denervated islets did not experience T2DM, indicating the necessity for autonomic innervation in the pathogenesis of Brsk2 caused T2DM.

On the contrary, mice with Brsk2 deletion in β cells show no cephalic insulin release in either NCD-or HFD-conditions and protect from HFD-induced metabolic abnormities. Physiologically, cephalic insulin release is fast and transient in nature and contributes to avoid postprandial hypoglycemia via HGP before nutrient absorption. Increased Brsk2 protein changes the cephalic nature and translates into prolonged insulin release message at both preabsorptive and absorptive stages after feeding. Thus, Brsk2 establishes a link between PNS and β cells physiologically; and if increased Brsk2 protein can trigger insulin resistance and T2DM.

Preclinical evidence has showed that mild suppression of hyperinsulinemia treats obesity (Page & Johnson, 2018). This might benefit from diminishing the β-cell hypersecretion burden which jeopardizes the maturation processes of insulin granules and results in proinsulin release to impose insulin resistance. Reports have showed that serum level of proinsulin is an independent risk factor of human T2DM (Riahi, Israeli et al., 2018). Chronic hyper(pro)insulinemia may interrupt the portal vein afferents (Dicostanzo, Dardevet et al., 2006) and cause autonomic imbalances in diet-induced obese individuals (Vinik, Maser et al., 2011) and in Brsk2-overexpressing mice, with effects that are characterized by an increase in parasympathetic afferent tone. In agreement with our study, Carlos et al. have shown that a hepatic branch vagotomy and disruption of vagal afferent fibers prevented glucocorticoid-induced diabetes and hypertension (Bernal-Mizrachi, Xiaozhong et al., 2007). We also found that ganglionic blockage and vagotomy successfully recovered the refeeding hyperglycemia in Brsk2-overexpressing mice, in agreement with other literatures.

Previous studies have shown that the functions of Brsk2 and AMPK in β cells have opposite roles in the regulation of mTORC1 signaling (Nie, Han et al., 2013a). Brsk2 is mainly located in the membrane, where it forms AIS and KA1 tight structures in its C-terminal to bind to acidic phospholipids; Brsk2 therefore differs from AMPK both in location and function (Wu, Cheng et al., 2015). We identified 289 proteins interacted with Brsk2 in MIN6 cells, including previously reported Ppp2r2a and Map2 (Wu et al., 2015). Among them, Gnb2, Gnas, and Arrb2 are well-known to function with GPCRs (Kawai, Sun et al., 2020, Mancini, Bertrand et al., 2015), and we experimentally confirmed an interaction between Brsk2 and Arrb2 (Supporting Information Fig S7 F and G). The GPCRs have a strong regulatory role in insulin secretion, and Brsk2 may connect to GPCRs effects, including gut hormones (Nie et al., 2013b), neurotransmitters, and FFAs. How Brsk2 senses glucose is unclear. Our results indicate that Brsk2 regulates insulin secretion secondary to cellular glucose metabolism, since Brsk2 overexpression decreases GCK and GLUT1(2) expression in human islets and MIN6 cells and Brsk2 knockdown increases these expressions (Supporting Information Fig S7 H and I).

Downstream substrates of kinase Brsk2 were also included to discover a direct secretogogue from Brsk2 interacted proteins. Brsk2 is a member of the calcium/calmodulin-dependent protein kinase (CAMK)-like (CAMKL) family. Comprehensively searching phosphorylated sites discovered that Plaa was a strong candidate (NetPhos 3.1 server). A recent literature has reported that hypomorphic mutations of PLAA cause an infantile-lethal neurodysfunction syndrome with seizures due to disrupting neurotransmission (Hall, Nahorski et al., 2017). Since the processes of insulin secretion from β cells largely resemble that of neurotransmitters release from neurons, our results that Plaa knockdown decreased insulin secretion may result from the likelihood pathway in neurons. Alternatively, reports have shown that Plaa enhances phospholipase A2 catalytic activity to generate arachidonic acid, thus eliciting Ca^2+^ oscillations via G protein/phospholipase A2/arachidonic acid cascades (Bomalaski, Steiner et al., 1992, Calignano, Piomelli et al., 1991, Tsunoda & Owyang, 1994). Here, we confirmed Plaa was also attributed to parasympathetic outflow-induced arachidonic acid production and insulin secretion as a downstream molecule of Brsk2 under metabolic-stress condition. Nevertheless, detailed mechanisms including specific phosphorylated site of Plaa modulated by parasympathetic acetylcholine/Chrm3/Brsk2 pathway still require future investigation.

In summary, we define a critical role of Brsk2 linking PNS to β cells in the development of insulin resistance and diabetes in mouse models. Moreover, three Brsk2 variants associated with human T2DM are also disclosed, providing clinically relevant evidence. We also demonstrate that pharmacological inhibition of Brsk2 improves glucose metabolism in HFD mice. Therefore, Brsk2 is a potential therapeutic target for preventing and curing overnutrition-induced diabetes.

## MATERIALS and METHODS

### Human Study

Human subjects were drawn from Shanghai Nicheng Cohort Study. Detailed cohort study has been published before (Chen, Hou et al., 2018). The single nucleotide polymorphisms (SNPs) in BRSK2 gene region were genotyped by Infinium Asian Screening Array and Infinium Multi-Ethnic Global BeadChip (Illumina, Inc., San Diego, CA, USA), and SNPs passed through quality-control procedure (individual call rate >98% and approved with Hardy-Weinberg equilibrium) were further imputed according to 504 East-Asian subjects in 1000 genome project phase III, SNPs with R-squared value >0.4 and minor allele frequency >0.01 were further analysis. The statical analysis were conducted by PLINK and SAS software (version 9.4; SAS Institute, Cary, NC). Descriptive statistics were compared by t test or non-parametric test according to data distribution. Genetic association analyses were performed by logistic and multiple linear regression adjusting for age and gender. Odds ratios or β values were calculated according to minor allele. The study was approved by the Human Research Ethics Committee of Shanghai Jiao Tong University Affiliated Sixth People’s Hospital. Written informed consents were acquired from all subjects.

### Genetic mouse models and animal care

Brsk2^fl/fl^ and MIP1-Cre transgenic (Tg) C57BL/6J mice were purchased from Jackson Laboratory in America. Male MIP1-cre-ERT; Brsk2^fl/fl^ mice and Brsk2^fl/fl^ mice aged 3-4 weeks were injected intraperitoneally with Tamoxifen (30 mg/kg body weight dissolved in corn oil, once a day) for 5 consecutive days to generate mature β-cell specific Brsk2 deletion mice (βKO mice) and their littermate controls (Ctrls). Mice were feeding with high fat diet (HFD: 60% of kcal from fat, 20% of kcal from carbohydrate and 20% of kcal from protein, Research Diets) or normal chow diet for indicated months.

Inducible β-cell specific Brsk2 transgenic mice (MIP1-rtTA; Brsk2) were constructed by the Model Animal Research Center of Nanjing University (Nanjing, China) based on Tet-On system. Line20 (13 copies), Line42 (35 copies), and Line29 (42 copies) were inbreeded with C57BL/6J mice at least twice to obtain stable genetic offsprings. Mice aged 6-8 weeks were subjected to drinking water containing doxycycline (Dox) with a concentration of 0.04-2 mg/ml to induce Brsk2 overexpression in vivo.

Animal studies were performed according to guidelines established by the Research Animal Care Committee of Nanjing Medical University, China (Permit Number: IACUC-NJMU 1404075). Mice were housed at 23°C -25°C using a 12 h light/12 h dark cycle. Animals had ad libitum access to water, and food was only withdrawn if required for an experiment. All experiments have been performed in adult mice except capsaicin treatment at newborn pups (postnatal day3-5).

### Primary human islet culture

Human islets were provided from Tianjin First Central Hospital. The use of human islets was approved by the research ethics committee of Nanjing Medical University. The use of human islets was approved by the Research Ethics Committee of Tianjin First Central Hospital. Primary human islets were transported in room temperature and were cultured in CMRL-1066 medium (glucose: 5.5 mmol/L) containing 10% FBS, 100 units/mL penicillin, and 100 mg/mL streptomycin. The primary islets were incubated at 37°C in a suitable atmosphere containing 95% O_2_ and 5% CO_2_.

### Primary mouse islet isolation and cell culture

Murine islets were isolated as described previously (Zhu, You et al., 2013). Primary isolated islets were cultured in RPMI-1640 medium (glucose: 5.5 mmol/L) containing 10% FBS, 100 units/mL penicillin, and 100 mg/mL streptomycin. Mouse pancreatic β-cell line MIN6 cells passed between 21 and 35, were cultured in DMEM (Invitrogen, Grand Island, NY) with 15% FBS (Gibco, Burlington, Ontario, Canada), 100 U/ml penicillin, 100 μg/ml streptomycin, 10 mM HEPES, and 50 μM β-mercaptoethanol (Sigma-Aldrich, St. Louis, MO) (Miyazaki, Araki et al., 1990). HEK293T cells were cultured in DMEM with 10% FBS. The primary islets and cells were incubated at 37°C in a suitable atmosphere containing 95% O_2_ and 5% CO_2_.

### Islet perfusion

120 islets per group were incubated 1 hour at 37°C in Krebs-Ringer buffer (KRB) solution with 2 mM glucose. Then islets were collected in a syringe filter (Millex-GP; Millipore) for further perfusion. 37°C KRB solution with 2 mM glucose was perfused at 125 µL/min for 15 minutes to equilibrate, then the perfusate were collected per minute for another 6 minutes. After that, 37°C KRB solution with 20 mM glucose were perfused for 25 minutes and the perfusate were collected as previous. 7-12 min was determined as the first phase insulin secretion while 12-22 min was defined as the second phase of insulin release.

### Transfection, infection, and GSIS

For siRNA-based interference assays, MIN6 cells and primary mouse islets were transfected with si*Brsk2* and siNC for 48 h and treated with carbachol for indicated time. For adenovirus infection, human primary islets (1.0×10^7^ pfu/ml), mouse primary islets (1.0×10^7^ pfu/ml) and MIN6 cells (2.0×10^6^ pfu/ml) were infected with Ad-Brsk2 or Ad-GFP for 24 h. For glucose-stimulated insulin secretion (GSIS) assays, islets or MIN6 cells were incubated in KRB buffer (135 mM NaCl, 3.62 mM KCl, 0.48 mM MgSO_4_ ·7H_2_O and 1.53 mM CaCl2, 0.2% BSA) for 1 h, and supernatants with low glucose (2 mmol/l for MIN6 cells, 3.3 mmol/l for primary islets) incubation for 1 h, and then high glucose (20 mmol/l for MIN6 cells, 16.7 mmol/l for primary islets) incubation for another 1 h were collected for measurement of insulin secretion. Cellular, islet and pancreatic insulin contents were extracted using an acid-ethanol solution (0.15 M HCl in 75% ethanol in H_2_O) overnight at 4°C. Insulin levels in supernatants and insulin contents were measured by radioimmunoassay (RIA) as previously described (Zhu et al., 2013) and normalized with total cellular DNA levels or weight of pancreas.

### Metabolic Studies

Blood samples were collected from tail vein and levels of blood glucose were measured using a Glucometer Elite monitor (Abbott, Oxon, UK). Random blood glucose (RBG) was measured at 9:00 in the morning. Fasting-blood-glucose (FBG) was measured after overnight fasting. IPGTTs and OGTTs were performed by intraperitoneal (i.p.) injection and oral gavage (IG) of D-glucose (1∼3 g/kg) respectively after overnight fasting. ITTs were performed by i.p. injection of 0.9 units/kg insulin after 4 h of fasting. PTTs were performed by i.p. injection of 1.5 g/kg sodium pyruvate after 16 h fasting.

### Hyperglycemia clamps

Preparation of the mice was same with hyperinsulinemic-euglycemic clamp assay. The mice were anesthetized by isoflurane inhalation after overnight fasting. After an overnight fast, mice were subjected to a 2-hour hyperglycemia clamp assay with a continuous infusion of 50% glucose at an adjusted rate to sustain the blood glucose level at ∼400 mg/dL. Glucose infusion rate, blood glucose and serum insulin levels were recorded at 0, 2, 5, 10 and every 10 minutes until the 120 minutes. 0 -20 min was determined as the first phase insulin secretion while 20 -120 min was defined as the second phase of insulin release.

### Hyperinsulinemic-euglycemic clamps

The mice were anesthetized by isoflurane inhalation and the right internal jugular vein catheterization was performed in a sterile environment. Mice were allowed 5 days postsurgical recovery. Mice with less than 5% weight lost were subsequently studied. FBG and body weight of the mice were measured and carefully placed in a mouse fixator for a few minutes balance. Mice were equilibrated from t = -90 to 0 min after 4-6 hours fasting. [3-^3^H] glucose (3 μCi; Moravek, California, USA) was administered at t = −90 min, followed by a constant infusion of 0.05 μCi/min. After 90 min as a basal period (t = −90 to 0 min), blood samples were collected from tail vein for determination of plasma glucose concentration and basal glucose specific activity. Then the continuous of human insulin (Humulin; Novo Nordisk) infusion was started (t = 0 min) at a rate of 4 mU/kg/min to keep hyperinsulinemic condition with submaximal suppression of HGP to assess insulin sensitivity. At 0 min, the continuous infusion rate of [3-^3^H]-D-glucose tracer was increased to 0.15 μCi/min for minimization of the changes of glucose specific activity. At t = 75 min, 10 μCi 2-[^14^C] D-glucose (Moravek, California, USA) was administered into each mouse to measure glucose uptake. Blood samples were collected at 10 min intervals from tail vein, blood glucose concentrations were measured by a glucose meter. During the 120 min clamp, variant glucose was simultaneously infused to keep the blood glucose concentration stable (∼130 mg/dL). The glucose infusion rate (GIR) was recorded to assess insulin sensitivity. At the end of clamp, additional blood samples were collected to determine the plasma glucose concentration and glucose specific activity. Liver, muscle, and adipose tissues were collected for the determination of radioactivity. Serum and tissue radioactivity were measured and calculated according to Kim JK (Kim, 2009).

### Mouse islet transplantation

Primary islets from WT mice and Tg mice were isolated and cultured within 72 h as described above. Cell viability of primary islets were stained by Acridine Orange / Propidium Iodide (AO/PI) solution according to the instruction before transplantation. Recipient WT mice were anesthetized by intraperitoneal administration of 40 mg/kg pentobarbital sodium, and then transferred to biosafety cabinet to shave skin. After disinfection with iodine, longitudinal wound about 1 cm was cut and kidney was gently moved out. Approximately 200 islet equivalents (IEQ) handpicked mouse islets were transplanted beneath the kidney capsule. During transplantation, physiological saline solution was used to moisten the wound continuously. Mice were sent to SPF facility for further study.

### Histology and immunostaining

Pancreas samples were rinsed in cold PBS and fixed overnight in 4% (g/vol) paraformaldehyde. The samples were then processed and embedded in paraffin, and consecutive sections were obtained. All the embedding, slicing, and H&E staining was done by Servicebio Technology (Wuhan, China). After dewaxing and antigen retrieval, the pancreatic tissue sections were incubated with anti-insulin (1:100 for sc-7839; 1:1000 for GB13121), and anti-Brsk2 (Cell signaling, 1:100) and were revealed using fluorescent-conjugated secondary antibodies (Invitrogen, 1:350) for multiple labeling. The nuclei were stained with Hoechst 33342 (5 μg/ml) for 5 min. Images were captured and analyzed by a confocal laser scanning microscope (Olympus FV1200).

### Immunoblotting and co-IP analysis

Immunoblotting assay was performed as previously described (Zhu et al., 2013). In brief, cells and primary islets were lysed with ice-cold lysis buffer containing 50 mM Tris-HCl (pH 7.4), 1% NP-40, 150 mM NaCl, 1 mM EDTA, 1 mM phenylmethylsulfonyl fluoride, and Complete protease inhibitor (Roche). Protein concentration was measured with BCA assay (Thermo Fisher Scientific). Samples were loaded on SDS-PAGE gels and then transferred to PVDF membranes (Bio-Rad). Immunoblotting was performed in 1% BSA buffer (50 mM Tris HCl pH7.4,150 mM NaCl, 5 mM EDTA, 0.05% Tween-20) with the corresponding antibodies. For co-immunoprecipitation (co-IP) assay, protein lysates were prepared from HEK293T or MIN6 cells in a buffer containing 50 mM Tris-HCl (pH 7.5), 150 mM NaCl, 0.1% NP-40, and a mixture of protease inhibitors (Roche). After centrifugation, 10% supernatants were used as inputs, and the rest were immunoprecipitated with agarose A/G beads (Roche) and mouse anti-flag antibody (Sigma) at 4°C overnight. Beads were washed three times with lysis buffer for 2min and collected for immunoblots. Antibodies were listed in *SI Appendix*, Table S5.

### Generation of stable overexpression MIN6 cell lines

The stable overexpressions of Brsk2 (WT) and Brsk2 (k48m) were provided by professor Yuguang Shi (UTHSCSA). In brief, MIN6 cells were transfected with pcDNA3.1-Brsk2 (WT), pcDNA3.1-Brsk2 (k48m), or pcDNA3.1 vector (Ctrl) plasmids for 36 h, and then subjected to G418 (200 μg/ml) selection for two days, followed with G418 (50 μg/ml) for another three days. The survived transfected MIN6 cells were passaged and maintained in culture medium with G418 (50 μg/ml) for two months.

### Plasmid construction and adenoviruses

The human Brsk2 overexpression plasmids pcDNA3.1-Brsk2 (WT) and kinase mutant pcDNA3.1-Brsk2 (k48m) as well as Brsk2 and control GFP adenoviruses were provided by Prof. Shi (Nie et al., 2013b). Ad-Brsk2 and Ad-GFP adenoviruses were amplified and purified by Hanbio biotechnology (Shanghai, China). pEGFP-Brsk2 plasmid was constructed by PCR amplified Brsk2 sequence from pcDNA3.1-Brsk2 (WT) and subcloned into the XhoI and EcoRI sites of pEGFP-C1 vector.

### In vivo autonomic antagonist administration

Dox-administrated mice were fasted overnight. Fasting blood glucose of the mice was measured before i.p. administrations of Atropine (10 mg/kg), Darifenacin hydrobromide (10 mg/kg), Hexamethonium bromide (10 mg/kg) or vehicle control respectively according to body weight. Mice were got access to food half hour post autonomic antagonists administration. Blood glucose was monitored at indicated times.

### Subdiaphragmatic vagotomy

Mice were fasted overnight and anesthetized by isoflurane inhalation. The subdiaphragmatic vagotomy was performed under a stereomicroscope with a magnification of 20 times. A 10-to 15-mm incision was made to expose esophagus and stomach in the peritoneal cavity. Both anterior and posterior vagal trunk were separated and a 2-mm section from each trunk was cut and removed to inhibit reinnervation. No attempt was made to perform a more selective vagotomy due to the size of the mice. Sham-operated mice underwent the same procedures except vagal nerves were kept intact. After the operation, the mice recovered for 5 days and then were exposed to Dox. Blood glucose, body weight, food intake and defecation of the mice were monitored three days post Dox-administration. When sacrificed, vagotomized mice showed dilated gastric ventricles which was not observed in sham-operated mice.

### Protein kinase assay

HEK293T cells were infected with Ad-GFP or Ad-Brsk2 for 24 h and the Flag-Brsk2 protein was then purified by immunoprecipitation using anti-flag antibody-conjugated beads. In vitro kinase reaction was performed in the presence of [γ-^32^P]-ATP and human C-terminal of Plaa protein (His-Plaa-CT, 5 µg), with or without 0.5 µg purified Flag-Brsk2. After incubation at 30 °C for 30 min, the kinase reaction was stopped by adding SDS loading buffer and samples were prepared for immunoblotting. Phosphorylation of Plaa was detected by the Amersham Typhoon 2.0.0.6 via autoradiography. Total protein levels of Flag-Brsk2 and His-Plaa-CT in kinase reactions were shown by ponceau S staining.

### Confocal live-cell imaging

MIN6 cells were plated in 3.5 cm glass bottom dishes and transfected with enhanced green fluorescent protein (EGFP)-tagged Brsk2. During imaging, cells were cultured at 37°C with 5% CO_2_ in a heated, gas-perfused chamber (Tokai engineering) and visualized with a confocal laser scanning microscope (Olympus FV1200). Cells were incubated with culture medium for 10 min to reach a stable statement in the chamber. One exclusive cell per dish was selected for recording according to the fluorescent intensity and cellular status. An image was recorded as time 0 and then stimulant drugs were added in the culture medium. Images were recorded at one-minute interval for 20 min.

### Transmission electron microscopy

Purified pancreatic islets were fixed with 2.5% glutaraldehyde in 0.1 mol/L sodium cacodylate buffer for 2 h and post-fixed in 1% OsO_4_, 1.5% K_4_Fe(CN)_6_, and 0.1 mol/L sodium cacodylate for 1 h. Islets were *en bloc* stained, dehydrated, embedded, and cut into ultrathin sections (50–80 nm). The samples were visualized by a Tecnai Spirit Biotwin operated at 200kV (FEI Company, Oregon, USA). At least fifty β cells were pictured for analysis.

### Metabolic cage analysis

Mice were induced with Dox for 2 months, and individually placed in a sterile metabolic cage unit (TSE Systems) for a multiday (5 days) study starting at 7 AM on day 1. Standard 12 h light (7 AM to 7 PM) and dark cycles (7 PM to 7 AM) were maintained throughout the experiment. Mice allowed to eat and drink *ad libitum*. Consumption of oxygen, exhalation of carbon dioxide, respiratory exchange rate, and activities were calculated and adjusted to body weight.

### Quantitative RT-PCR analysis

Total RNA was extracted using TRIzol reagent (Invitrogen). The synthesis of the cDNA was carried out using the HiScript II Q RT SuperMix (Vazyme). Quantitative PCR (qPCR) assay was performed by using the SYBR qPCR Master Mix (Vazyme) on Roche Lightcycle480 II Sequence Detection System (Roche). Gene expression levels were normalized by *Actb*. Primer sequences for qPCR are listed in *SI Appendix* Table S5.

### RNA-sequencing analysis

Mice were induced with Dox for 4 days when primary islets were isolated for total RNA extraction. A total amount of 3 µg RNA per sample was used as input material for the RNA sample preparations. Sequencing libraries were generated using NEBNext® UltraTM RNA Library Prep Kit for Illumina® (NEB, USA) following manufacturer’s recommendations and index codes were added to attribute sequences to each sample. Clustering, sequencing and data analysis were done by Novogene (Tianjin, China). In brief, sequencing analysis was done using mRNA-seq analysis on Maverix Analytic Platform (Maverix Biomics, San Mateo, CA). Single reads were mapped to the mouse genome using STAR in a strand specific manner. Pairwise differential expression was quantified using Cuffdiff. Cufflinks was used to determine FPKM levels for each gene from the STAR alignment and was used as input for Cuffdiff. Read counts were then normalized across all samples and significant differentially expressed genes were determined by adjusted *P* value with a threshold of 0.05. Gene Set Enrichment Analysis (GSEA) according to all sequenced genes were performed for identification the biological processes, molecular function, and cellular component involved in Brsk2 mediated effects (Novogene). RNA sequencing data were available in GEO: GSE151915, and GSE179893.

### Statistical analysis

In vitro experiments were repeated at least three times and in vivo assay were repeated two times, with the number of per condition or mice included in each group in each experiment indicated. Comparisons were performed using the Student t test between two groups or ANOVA in multiple groups. Results are presented as mean ± SEM. P < 0.05 is considered statistically significant.

## Acknowledgments

We thank Professor Yuguang Shi (Barshop Institute for Longevity and Aging Studies, Depart of Pharmacology, University of Texas Health Science Center at San Antonio, San Antonio, Texas 78229) for providing Brsk2 plasmids, adenoviruses and stable MIN6 cell lines. This study was supported by research grants from the National Natural Science Foundation of China (81420108007 and 81830024 to X.H.; 81670703, 81870531 and 82070843 to Y-X.Z.) and the National Key Research and Development Program of China (2016YFC1304804) to X.H. X.H. and Y-X.Z. are fellows at the Collaborative Innovation Center for Cardiovascular Disease Translational Medicine.

## Author Contributions

Conceptualization, X.H., Y-X.Z. and C.H.; Methodology, Y-X.Z., R.X., K-Y.W., Z.Y., L.J., J.P., Y-C.Z., K.W., D.L., Y-Q.Z., P.S., and Y.L.; Formal Analysis, Y-X.Z., R.X., Y-L.Z., and S.L.; Investigation, Y-X.Z., R.X., K-Y.W., Z.Y., L.J., J.P., K.W. and D.L.; Resources, F.W., X.C., S.W. and C.H.; Writing -Original Draft, Y-X.Z. and C.H.; Writing – Review & Editing, X.H., Y-X.Z., Y-L.Z., and S.L.; Visualization, Y-X.Z.; Supervision, X.H., Y-X.Z. and C.H.; Funding Acquisition, X.H., and Y-X.Z. All authors reviewed and commended on the manuscript.

## Competing Interest Statement

The authors declare that they have no competing interests.

## References

Ahren B (2000) Autonomic regulation of islet hormone secretion--implications for health and disease. Diabetologia 43: 393–410

Ahren B, Simonsson E, Scheurink AJ, Mulder H, Myrsen U, Sundler F (1997) Dissociated insulinotropic sensitivity to glucose and carbachol in high-fat diet-induced insulin resistance in C57BL/6J mice. Metabolism: clinical and experimental 46: 97–106

Alarcon C, Boland BB, Uchizono Y, Moore PC, Peterson B, Rajan S, Rhodes OS, Noske AB, Haataja L, Arvan P, Marsh BJ, Austin J, Rhodes CJ (2016) Pancreatic beta-Cell Adaptive Plasticity in Obesity Increases Insulin Production but Adversely Affects Secretory Function. Diabetes 65: 438–50

Alvarsson A, Jimenez-Gonzalez M, Li R, Rosselot C, Tzavaras N, Wu Z, Stewart AF, Garcia-Ocana A, Stanley SA (2020) A 3D atlas of the dynamic and regional variation of pancreatic innervation in diabetes. Sci Adv 6

Berger M, Scheel DW, Macias H, Miyatsuka T, Kim H, Hoang P, Ku GM, Honig G, Liou A, Tang Y, Regard JB, Sharifnia P, Yu L, Wang J, Coughlin SR, Conklin BR, Deneris ES, Tecott LH, German MS (2015) Galphai/o-coupled receptor signaling restricts pancreatic beta-cell expansion. Proceedings of the National Academy of Sciences of the United States of America 112: 2888–93

Bernal-Mizrachi C, Xiaozhong L, Yin L, Knutsen RH, Howard MJ, Arends JJ, Desantis P, Coleman T, Semenkovich CF (2007) An afferent vagal nerve pathway links hepatic PPARalpha activation to glucocorticoid-induced insulin resistance and hypertension. Cell metabolism 5: 91–102

Berthoud HR, Albaugh VL, Neuhuber WL (2021) Gut-brain communication and obesity: understanding functions of the vagus nerve. The Journal of clinical investigation 131

Bomalaski JS, Steiner MR, Simon PL, Clark MA (1992) IL-1 increases phospholipase A2 activity, expression of phospholipase A2-activating protein, and release of linoleic acid from the murine T helper cell line EL-4. Journal of immunology 148: 155–60

Boyle PJ, Liggett SB, Shah SD, Cryer PE (1988) Direct muscarinic cholinergic inhibition of hepatic glucose production in humans. The Journal of clinical investigation 82: 445–9

Brandt C, Nolte H, Henschke S, Engstrom Ruud L, Awazawa M, Morgan DA, Gabel P, Sprenger HG, Hess ME, Gunther S, Langer T, Rahmouni K, Fenselau H, Kruger M, Bruning JC (2018) Food Perception Primes Hepatic ER Homeostasis via Melanocortin-Dependent Control of mTOR Activation. Cell 175: 1321–1335 e20

Burris RE, Hebrok M (2007) Pancreatic innervation in mouse development and beta-cell regeneration. Neuroscience 150: 592–602

Calignano A, Piomelli D, Sacktor TC, Schwartz JH (1991) A phospholipase A2-stimulating protein regulated by protein kinase C in Aplysia neurons. Brain research Molecular brain research 9: 347–51

Chen P, Hou X, Hu G, Wei L, Jiao L, Wang H, Chen S, Wu J, Bao Y, Jia W (2018) Abdominal subcutaneous adipose tissue: a favorable adipose depot for diabetes? Cardiovasc Diabetol 17: 93

Corrall RJ, Frier BM (1981) Acute hypoglycemia in man: neural control of pancreatic islet cell function. Metabolism: clinical and experimental 30: 160–4

Cryer PE (2004) Diverse causes of hypoglycemia-associated autonomic failure in diabetes. N Engl J Med 350: 2272–9

Cryer PE (2006) Mechanisms of sympathoadrenal failure and hypoglycemia in diabetes. The Journal of clinical investigation 116: 1470–3

Dicostanzo CA, Dardevet DP, Neal DW, Lautz M, Allen E, Snead W, Cherrington AD (2006) Role of the hepatic sympathetic nerves in the regulation of net hepatic glucose uptake and the mediation of the portal glucose signal. American journal of physiology Endocrinology and metabolism 290: E9–E16

Franks PW, McCarthy MI (2016) Exposing the exposures responsible for type 2 diabetes and obesity. Science 354: 69–73

Gautam D, Han SJ, Hamdan FF, Jeon J, Li B, Li JH, Cui Y, Mears D, Lu H, Deng C, Heard T, Wess J (2006) A critical role for beta cell M3 muscarinic acetylcholine receptors in regulating insulin release and blood glucose homeostasis in vivo. Cell metabolism 3: 449–61

Gromada J, Chabosseau P, Rutter GA (2018) The alpha-cell in diabetes mellitus. Nature reviews Endocrinology 14: 694–704

Grzmil M, Huber RM, Hess D, Frank S, Hynx D, Moncayo G, Klein D, Merlo A, Hemmings BA (2014) MNK1 pathway activity maintains protein synthesis in rapalog-treated gliomas. The Journal of clinical investigation 124: 742–54

Guo Z, Tang W, Yuan J, Chen X, Wan B, Gu X, Luo K, Wang Y, Yu L (2006) BRSK2 is activated by cyclic AMP-dependent protein kinase A through phosphorylation at Thr260. Biochemical and biophysical research communications 347: 867–71

Hall EA, Nahorski MS, Murray LM, Shaheen R, Perkins E, Dissanayake KN, Kristaryanto Y, Jones RA, Vogt J, Rivagorda M, Handley MT, Mali GR, Quidwai T, Soares DC, Keighren MA, McKie L, Mort RL, Gammoh N, Garcia-Munoz A, Davey T et al. (2017) PLAA Mutations Cause a Lethal Infantile Epileptic Encephalopathy by Disrupting Ubiquitin-Mediated Endolysosomal Degradation of Synaptic Proteins. American journal of human genetics 100: 706–724

Inoguchi T, Li P, Umeda F, Yu HY, Kakimoto M, Imamura M, Aoki T, Etoh T, Hashimoto T, Naruse M, Sano H, Utsumi H, Nawata H (2000) High glucose level and free fatty acid stimulate reactive oxygen species production through protein kinase C--dependent activation of NAD(P)H oxidase in cultured vascular cells. Diabetes 49: 1939–45

Kahn SE, Hull RL, Utzschneider KM (2006) Mechanisms linking obesity to insulin resistance and type 2 diabetes. Nature 444: 840–6

Kawai T, Sun B, Yoshino H, Feng D, Suzuki Y, Fukazawa M, Nagao S, Wainscott DB, Showalter AD, Droz BA, Kobilka TS, Coghlan MP, Willard FS, Kawabe Y, Kobilka BK, Sloop KW (2020) Structural basis for GLP-1 receptor activation by LY3502970, an orally active nonpeptide agonist. Proceedings of the National Academy of Sciences of the United States of America 117: 29959–29967

Kim JK (2009) Hyperinsulinemic-euglycemic clamp to assess insulin sensitivity in vivo. Methods in molecular biology 560: 221–38

Liggett SB (2009) alpha2A-adrenergic receptors in the genetics, pathogenesis, and treatment of type 2 diabetes. Sci Transl Med 1: 12ps15

Lilley BN, Pan YA, Sanes JR (2013) SAD kinases sculpt axonal arbors of sensory neurons through long-and short-term responses to neurotrophin signals. Neuron 79: 39–53

Magnan C, Collins S, Berthault MF, Kassis N, Vincent M, Gilbert M, Penicaud L, Ktorza A, Assimacopoulos-Jeannet F (1999) Lipid infusion lowers sympathetic nervous activity and leads to increased beta-cell responsiveness to glucose. The Journal of clinical investigation 103: 413–9

Mancini AD, Bertrand G, Vivot K, Carpentier E, Tremblay C, Ghislain J, Bouvier M, Poitout V (2015) beta-Arrestin Recruitment and Biased Agonism at Free Fatty Acid Receptor 1. The Journal of biological chemistry 290: 21131–40

Meek TH, Nelson JT, Matsen ME, Dorfman MD, Guyenet SJ, Damian V, Allison MB, Scarlett JM, Nguyen HT, Thaler JP, Olson DP, Myers MG, Jr., Schwartz MW, Morton GJ (2016) Functional identification of a neurocircuit regulating blood glucose. Proceedings of the National Academy of Sciences of the United States of America 113: E2073–82

Miyazaki J, Araki K, Yamato E, Ikegami H, Asano T, Shibasaki Y, Oka Y, Yamamura K (1990) Establishment of a pancreatic beta cell line that retains glucose-inducible insulin secretion: special reference to expression of glucose transporter isoforms. Endocrinology 127: 126–32

Nie J, Han X, Shi Y (2013a) SAD-A and AMPK kinases: the “yin and yang” regulators of mTORC1 signaling in pancreatic beta cells. Cell cycle 12: 3366–9

Nie J, Lilley BN, Pan YA, Faruque O, Liu X, Zhang W, Sanes JR, Han X, Shi Y (2013b) SAD-A potentiates glucose-stimulated insulin secretion as a mediator of glucagon-like peptide 1 response in pancreatic beta cells. Molecular and cellular biology 33: 2527–34

Nie J, Liu X, Lilley BN, Zhang H, Pan YA, Kimball SR, Zhang J, Zhang W, Wang L, Jefferson LS, Sanes JR, Han X, Shi Y (2013c) SAD-A kinase controls islet beta-cell size and function as a mediator of mTORC1 signaling. Proceedings of the National Academy of Sciences of the United States of America 110: 13857–62

Nie J, Sun C, Faruque O, Ye G, Li J, Liang Q, Chang Z, Yang W, Han X, Shi Y (2012) Synapses of amphids defective (SAD-A) kinase promotes glucose-stimulated insulin secretion through activation of p21-activated kinase (PAK1) in pancreatic beta-Cells. The Journal of biological chemistry 287: 26435–44

O’Brien PD, Hinder LM, Callaghan BC, Feldman EL (2017) Neurological consequences of obesity. The Lancet Neurology 16: 465–477

Oancea E, Bezzerides VJ, Greka A, Clapham DE (2003) Mechanism of persistent protein kinase D1 translocation and activation. Developmental cell 4: 561–74

Page MM, Johnson JD (2018) Mild Suppression of Hyperinsulinemia to Treat Obesity and Insulin Resistance. Trends in endocrinology and metabolism: TEM 29: 389–399

Paschen M, Moede T, Valladolid-Acebes I, Leibiger B, Moruzzi N, Jacob S, Garcia-Prieto CF, Brismar K, Leibiger IB, Berggren PO (2019) Diet-induced beta-cell insulin resistance results in reversible loss of functional beta-cell mass. FASEB journal : official publication of the Federation of American Societies for Experimental Biology 33: 204–218

Razavi R, Chan Y, Afifiyan FN, Liu XJ, Wan X, Yantha J, Tsui H, Tang L, Tsai S, Santamaria P, Driver JP, Serreze D, Salter MW, Dosch HM (2006) TRPV1+ sensory neurons control beta cell stress and islet inflammation in autoimmune diabetes. Cell 127: 1123–35

Remedi MS, Emfinger C (2016) Pancreatic beta-cell identity in diabetes. Diabetes, obesity & metabolism 18 Suppl 1: 110–6

Riahi Y, Israeli T, Cerasi E, Leibowitz G (2018) Effects of proinsulin misfolding on beta-cell dynamics, differentiation and function in diabetes. Diabetes, obesity & metabolism 20 Suppl 2: 95–103

Roden M, Shulman GI (2019) The integrative biology of type 2 diabetes. Nature 576: 51–60

Rodriguez-Diaz R, Caicedo A (2014) Neural control of the endocrine pancreas. Best practice & research Clinical endocrinology & metabolism 28: 745–56

Rosengren AH, Jokubka R, Tojjar D, Granhall C, Hansson O, Li DQ, Nagaraj V, Reinbothe TM, Tuncel J, Eliasson L, Groop L, Rorsman P, Salehi A, Lyssenko V, Luthman H, Renstrom E (2010) Overexpression of alpha2A-adrenergic receptors contributes to type 2 diabetes. Science 327: 217–20

Sumara G, Formentini I, Collins S, Sumara I, Windak R, Bodenmiller B, Ramracheya R, Caille D, Jiang H, Platt KA, Meda P, Aebersold R, Rorsman P, Ricci R (2009) Regulation of PKD by the MAPK p38delta in insulin secretion and glucose homeostasis. Cell 136: 235–48

Tang SC, Baeyens L, Shen CN, Peng SJ, Chien HJ, Scheel DW, Chamberlain CE, German MS (2018) Human pancreatic neuro-insular network in health and fatty infiltration. Diabetologia 61: 168–181

Tarussio D, Metref S, Seyer P, Mounien L, Vallois D, Magnan C, Foretz M, Thorens B (2014) Nervous glucose sensing regulates postnatal beta cell proliferation and glucose homeostasis. The Journal of clinical investigation 124: 413–24

Teff KL (2011) How neural mediation of anticipatory and compensatory insulin release helps us tolerate food. Physiology & behavior 103: 44–50

Tsunoda Y, Owyang C (1994) A newly cloned phospholipase A2-activating protein elicits Ca2+ oscillations and pancreatic amylase secretion via mediation of G protein beta/phospholipase A2/arachidonic acid cascades. Biochemical and biophysical research communications 203: 1716–24

Vinik AI, Maser RE, Ziegler D (2011) Autonomic imbalance: prophet of doom or scope for hope? Diabetic medicine : a journal of the British Diabetic Association 28: 643–51

Wang K, Liu D, Zhang Y, Chang X, Xu R, Pang J, Li K, Sun P, Zhu Y, Han X (2019) SAD-A, a downstream mediator of GLP-1 signaling, promotes the phosphorylation of Bad S155 to regulate in vitro beta-cell functions. Biochemical and biophysical research communications 509: 76–81

Winzell MS, Ahren B (2007) G-protein-coupled receptors and islet function-implications for treatment of type 2 diabetes. Pharmacology & therapeutics 116: 437–48

Wu JX, Cheng YS, Wang J, Chen L, Ding M, Wu JW (2015) Structural insight into the mechanism of synergistic autoinhibition of SAD kinases. Nature communications 6: 8953

Zhu Y, You W, Wang H, Li Y, Qiao N, Shi Y, Zhang C, Bleich D, Han X (2013) MicroRNA-24/MODY gene regulatory pathway mediates pancreatic beta-cell dysfunction. Diabetes 62: 3194–206

